# Exploring the role of local sequence context on patterns of germline mutation

**DOI:** 10.64898/2026.10.20.760577

**Authors:** Andrew T. Beck, Jun Z. Li, Sebastian Zoellner

## Abstract

Germline mutations occur at a non-uniform rate across the genome. While the influence of neighboring nucleotides is well known and accounted for in mutation models, the effective range of these flanking influences and the extent of their higher-order interactions remain insufficiently understood.

Here, we leverage millions of singleton variants from the 1000 Genomes Project as a proxy for *de novo* variants to systematically map the contextual influence of local nucleotide sequence context on germline mutation rates. We show that while nucleotides immediately adjacent to the mutation site exert the strongest effect, the influence of nucleotides at flanking positions does not decay monotonically with increased distance from the focal position. Notably, significant associations between flanking nucleotides and mutation status persist up to hundreds of base pairs from the focal site. Furthermore, higher-order interactions among flanking positions significantly modulate mutation rates, though their effect size diminishes as the order of interaction increases. Additionally, these mutational signatures are highly consistent across diverse ancestral groups. Together, these findings provide a comprehensive map of how local genomic architecture shapes the mutational landscape.

## 1 Introduction

Germline mutagenesis is the primary source of disease causing alleles and genetic variation upon which evolutionary forces such as natural selection act. Consequently, quantifying the rate and distribution of mutations is central to our understanding of evolutionary biology and disease risk [1, 2]. In humans, germline mutagenesis results in an average mutation rate of 1 − 1.5× 10^−8^ per base pair per generation [3]. However, this rate is not uniform across the genome, and it is influenced by a complex landscape of genomic and epigenomic features, including recombination rate [4], replication timing [5], and epigenetic markers [6]. Among these, local nucleotide sequence context, the neighboring nucleotide bases flanking a site, is a major determinant of mutational variability [7–10] as it directly influences local DNA physical structure, chemical stability, and the binding dynamics of replication and repair machinery. Fully characterizing how these and other genomic features influence mutation rates is essential to understanding the mechanisms that drive genetic diversity and human evolution.

To account for the influence of nucleotides at flanking positions, mutational models typically parameterize sequence context by classifying variants into *k*-mer motifs centered at the mutation site [7, 9, 10]. This approach estimates distinct mutation rates for each motif, capturing the combined influence of neighboring nucleotides within the window. For example, Carlson et al. [9] demonstrated that mutation rate heterogeneity depends primarily on adjacent nucleotides and that the effects of broader genomic features often vary in both magnitude and direction across different sequence contexts. While expanding the *k*-mer window size captures more of this heterogeneity, the number of possible motifs increases exponentially with window length (4^*k*^), typically limiting models to 7-mers (± 3 bp). To circumvent this exponential scaling, an alternative approach is to add independent parameters for more distant flanking positions [10]. However, this additive strategy only captures the marginal effects of distant positions, completely ignoring potential higher-order interactions between them. Consequently, determining whether it is sufficient to model only first-order effects at extended flanking distances, or if complex interactions persist further out, requires a more thorough profiling of the influence of local sequence context on rates of germline mutation.

Local sequence context also interacts with broader genomic and epigenomic features to shape germline mutation rates [9], meaning these confounding factors must be controlled to isolate the precise role of flanking nucleotides. Because direct, experimental measurement of mutational processes in human germline tissue remains challenging, researchers rely on the genomic distribution of observed variants as a empirical readout of mutational activity. However, evaluation of these distributions while also accounting for genomic confounders is not straightforward; for example, histone modifications such as H3K36me, which are known to be associated with mutation rates, might only be available from somatic cell types not directly associated with germline mutagenesis. To address this, Zhu et al. [8] proposed a location-matched control strategy, pairing each variant with a nearby non-variant site. This approach assumes that neighboring sites share a nearly identical genomic and epigenomic environment, allowing the specific influence of flanking nucleotides to be isolated. While Zhu et al. [8] found that the influence of flanking positions and their interactions decays with distance, their analysis was largely restricted to a ±4 bp window, with individual effects considered only up to 10 bp. Furthermore, their reliance on common non-coding variants introduces confounding from long-term natural selection and biased gene conversion, forces that actively distort the baseline mutational signature. To capture the true biochemical footprint of germline mutation processes active in modern human populations, it is instead necessary to evaluate evolutionarily newer variants, whose distribution across the genome is less impacted by these forces.

To overcome the limitations inherent to common variants, we adapt the location-matched control method of Zhu et al. [8] to a large-scale dataset of singleton variants from the 1000 Genomes Project (1kGP) 30x deep-sequencing release [11]. Because singletons represent evolutionarily recent events, they are minimally influenced by natural selection or biased gene conversion, thereby providing a clearer window into contemporary germline mutagenesis [9]. Analyzing 2,504 unrelated individuals across five global super-populations, we quantify marginal effects at flanking positions up to 1,000 bp from the focal site and evaluate higher-order interactions within a ±10 bp window. We demonstrate that statistically significant associations between flanking alleles and mutation status persist well beyond the previously studied ± 10 bp range for all mutation subtypes. While the strongest marginal and two-way effects occur within ± 4 bp, this decay in influence is notably non-monotonic in several subtypes. Furthermore, while higher-order interactions are less influential than lower-order effects, they provide critical nuance to the mutational landscape. These patterns are largely subtype-specific yet remarkably consistent across diverse human populations, suggesting that the underlying mechanisms of local sequence dependency are shared.

## 2 Methods

### 2.1 Singletons and Matched Controls

To characterize the influence of flanking sequence context on germline mutation processes, we analyzed singleton single-nucleotide substitutions from 2,504 unrelated individuals in the 1000 Genomes Project 30x high-coverage release [11]. We restricted the variant set to autosomal singletons and private doubletons (variants observed twice but within a single individual) to minimize the confounding effects of natural selection and biased gene conversion. Mutations were categorized into six classes based on complementary symmetry (e.g., *A→G* and *T→C* were grouped). Given the distinct mutational dynamics associated with CpG dinucleotides [12], *C→N* mutations were further stratified into CpG and non-CpG contexts, resulting in a total of nine classes of mutations.

To isolate the effects of local sequence context from large-scale genomic and epigenomic features, we employed a location-matched control strategy. Each singleton was paired with five monomorphic control sites randomly sampled from the reference genome. These controls were required to have the same reference allele as the singleton and be located within a ± 150 bp window of the variant site. By selecting controls in such close proximity, we ensure that the variant and its corresponding null sites share nearly identical large-scale genomic environments, such as replication timing and recombination rate [8].

### 2.2 Log Linear Models for Single Flanking Position Influence

To quantify the influence of a single flanking position *k* ∈ {−1000, …, +1000} on mutation subtype *t*, we compared the distribution of nucleotides at that position between singletons and their matched controls. Let *S*_*t*_ and *C*_*t*_ denote the total number of singletons and matched controls, respectively, observed for subtype *t* across the genome. At each flanking position *k*, the counts of both the singletons and their matched controls for each subtype *t A*? *{G*, … *CpG* ?*TpG*} were stratified into vectors of allele counts 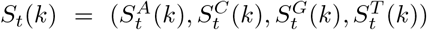 and 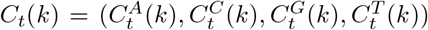, where 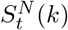 denotes the number of singletons of subtype *t* who are flanked by nucleotide *N* at position *k* relative to the variant site while 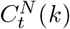 denotes the number of controls for subtype *t* flanked by nucleotide *N* at *k* positions relative to the control’s location. Under the hypothesis that nucleotides at flanking position *k* have no influence on mutation type *t*, we expect that 5 · *S*_*t*_(*k*) = *C*_*t*_(*k*) (since each singleton was matched to five non-variant controls) and the proportions *S*_*t*_(*k*)*/S*_*t*_ and *C*_*t*_(*k*)*/C*_*t*_ should be equivalent.

We tested this null hypothesis using log-linear models. Let 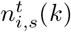 represent the count of observations for subtype *t* partitioned by singleton/control status *s* and nucleotide *i* at position *k*. We fit the following reduced model for each subtype *t* and flanking position *k*:

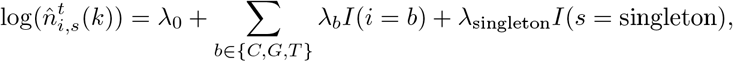

where *I* indicates the indicator function and the *λ* parameters represent regression weights. This model assumes no interaction between the nucleotide identity and the singleton/control status. To measure the deviation from this null assumption, we compared the reduced model to a fully saturated model that models impact of flanking genotype on mutation as interaction between mutation status and flanking position:

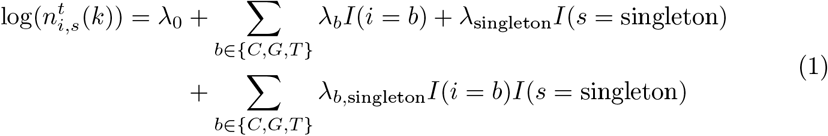

The fit of the reduced model relative to the saturated model was evaluated using the deviance statistic:

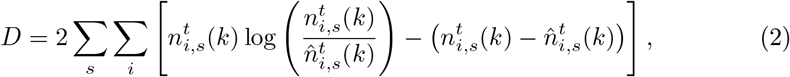

where 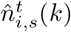 is the expected count under the reduced null model.

Under the null hypothesis,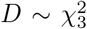. Because mutation subtypes and population groups vary significantly in sample size (*N*_*t*_), we normalized this statistic to obtain the relative entropy (*RE*), providing a scale-invariant measure of influence:

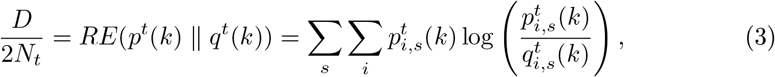

where 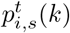 and 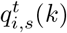 are the observed and expected joint proportions, respectively. This normalized metric allows for direct comparison of mutational influence across different genomic contexts and datasets.

### 2.3 Log-Linear Models for the Influence of Interactions

To assess the influence of two-way interactions between flanking nucleotides, we stratified singletons and controls by the nucleotide pair (*i, j*) ∈{*A, C, G, T}* ^2^ at positions (*k*_1_, *k*_2_). To isolate the specific contribution of the interaction to the mutation process, we fit a reduced log-linear model that models all lower-order effects but assumes the interaction between the two nucleotides does not vary by mutation status. This yields the null expected counts 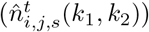:

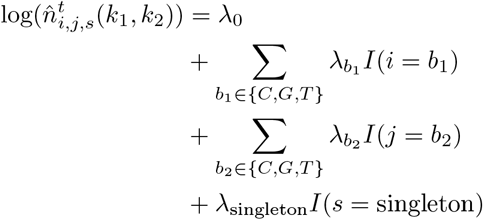

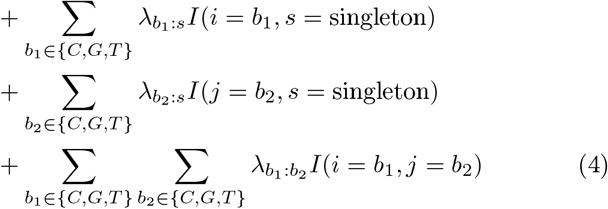

for all pairs of flanking positions (*k*_1_, *k*_2_) within a ± 10 bp window centered on the focal site. By including base-by-status interactions 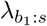 and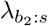, this model explicitly controls for the marginal differences in single-nucleotide distributions between singletons and controls identified in the previous section. Additionally, non-independence of the nucleotides at the two flanking positions (e.g., due to background genomic sequence composition) is modeled by a base-base interaction term 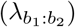. Critically, because this term is shared between singletons and their location-matched controls, it is independent of the mutation process.

We compared these expected counts against a fully saturated model that perfectly tracks the observed counts 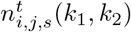 by introducing a three-way interaction term 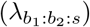, allowing the co-occurrence of the two nucleotides to differ conditionally upon mutation status:

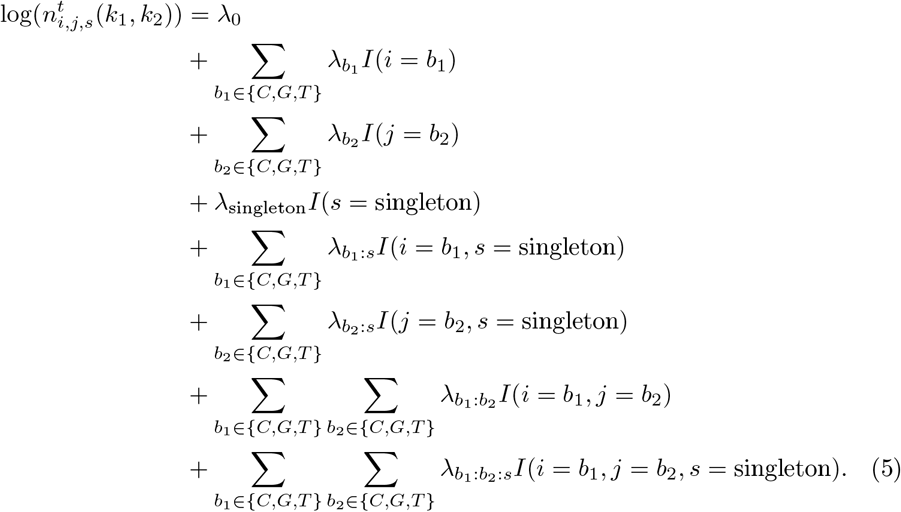

Under the null hypothesis that the interaction between positions *k*_1_ and *k*_2_ does not modulate the mutation rate, the deviance *D* calculated between these models asymptotically follows a chi-squared distribution with nine degrees of freedom 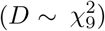. As with the marginal effects, we normalized the deviance by 2*N*_*t*_ to obtain a scale-invariant relative entropy measure, enabling direct comparisons across mutation subtypes and populations.

To evaluate higher-order genomic dependencies, we generalized this nested log-linear framework to multi-way interactions. For an interaction of order *m* (involving *m* flanking positions), the reduced model incorporates all marginal effects and lower-order interactions up to order *m* − 1, alongside their respective interaction terms with the sequence status *s*. The unique influence of the *m*-way interaction is isolated by comparing this reduced model to a saturated model containing the (*m* + 1)-way interaction term between the *m* nucleotides and status *s*. For example, a three-way nucleotide interaction (*m* = 3) yields a deviance statistic that asymptotically follows a 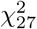 distribution (3^3^ degrees of freedom). Due to the high coverage of the 1kGP

30x dataset and our large sample of singletons, cell counts remained sufficiently high to support these higher-dimensional contingency tables without suffering from severe sparsity, ensuring robust and consistent estimation of relative entropy across all tested interaction orders.

## 3 Results

To more thoroughly characterize the influence of local sequence context on mutation rates, we first quantified the influence of individual flanking nucleotides on germline mutation rates within a window of ± 1, 000 bp using our genomic location-matched design. We then evaluated the synergistic effects of nucleotides at different flanking positions by measuring second- and third-order interactions among nucleotides at positions within a ± 10 bp radius of the focal mutation site. To assess whether these local sequence dependencies reflect universal patterns or population-specific traits, we analyzed autosomal singletons both globally across the aggregated 1000 Genomes Project (1kGP) dataset and stratified across the five global super-populations.

### 3.1 Mutational influence of individual flanking nucleotides within *±*1, 000 bp window

We quantified the marginal effect of sequence context at each flanking position across all mutation subtypes, accounting for the distinct mutational dynamics of CpG dinucleotides by stratifying all *C→N* changes into CpG and non-CpG subtypes. Across all mutation classes, we observed highly statistically significant deviance within the ± 10 bp window (deviance range: 18.6 to 1, 659, 567; *χ*^2^ df = 3, *p* ≤ 3.24 ×10^−4^; Supplementary Table S1). Because average deviance was positively correlated with subtype sample size (Pearson *r* = 0.765, *p* = 0.02), we normalized the deviance statistics by twice the sample size to obtain relative entropy (*RE*) values, enabling cross-subtype comparisons (Figure 1). Among the non-CpG mutation types, the highest observed influence occurred at the +1 position for *A→G* mutations (*RE* = 9.6 ×10^−3^), mirroring previous observations derived from common non-coding variants [8]. To contextualize the magnitude of these *RE* statistic, had we not considered CpG mutation subtypes as their own categories, a *RE* statistic of 0.0554 would have been observed at +1 for the *C*? *T* subtype, reflecting the extreme hypermutability of methylated CpG sites. While the strongest influence typically occurred at the immediate neighbors (± 1 bp), we identified notable exceptions; for example, the *C→G* subtype exhibited its peak influence at the − 2 position. Overall, *RE* values decreased as a function of distance from the focal site, though this decay was non-monotonic and the relative importance of specific flanking positions remained highly subtype-dependent.

**Fig. 1:**
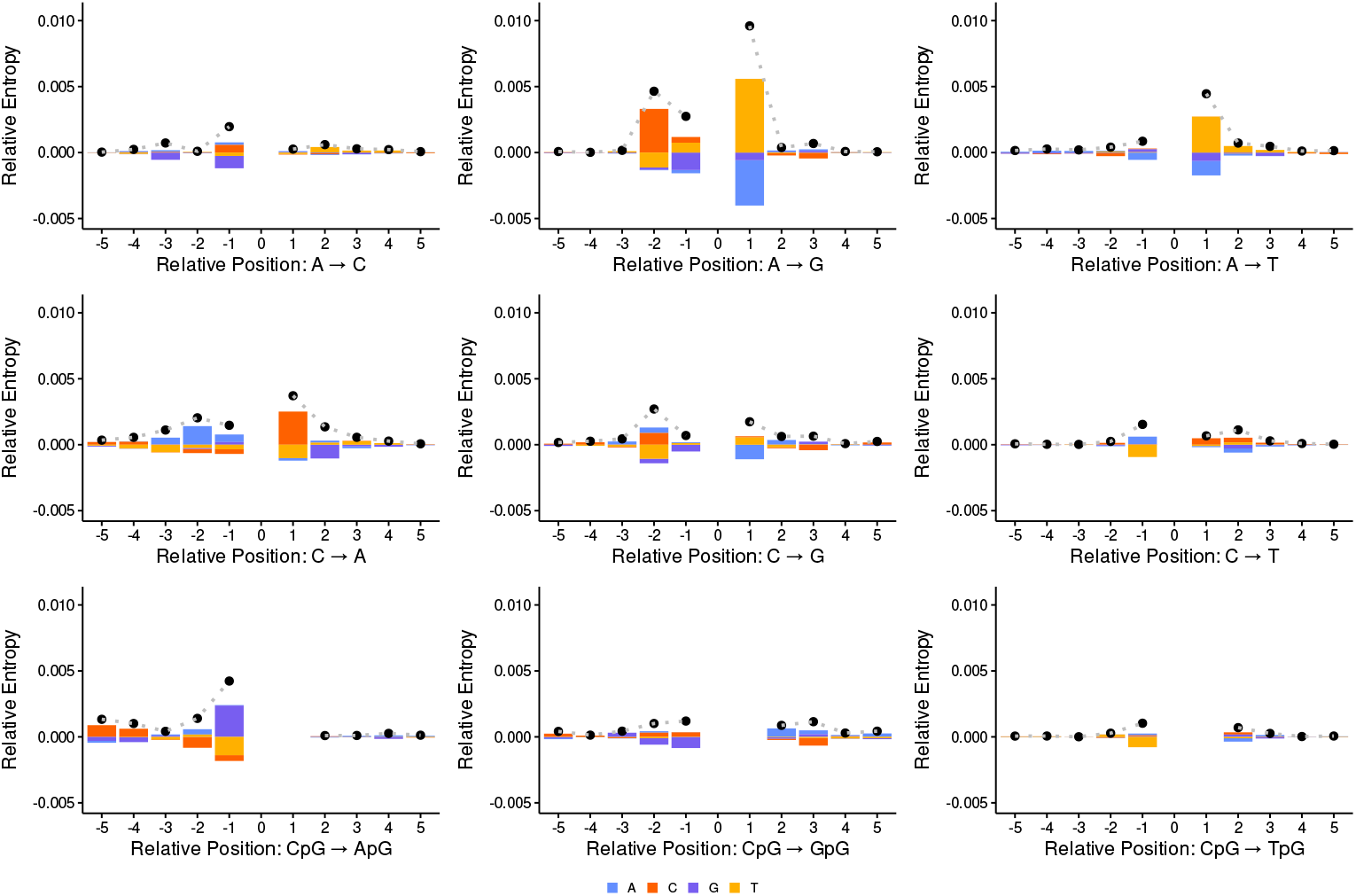
Single-position mutational influence and nucleotide-level residuals across nine mutation subtypes within a 10 bp window. For each subtype, the overall flanking influence at each position *k* ±∈{−10, …, +10} is quantified by relative entropy (*RE*, represented by the scatterplot points). The underlying bars at each position display the signed log-linear model residuals for each nucleotide (*A, C, G, T*), where positive values denote enrichment and negative values denote depletion in singletons relative to the baseline genomic controls. Note that *C* ?*N* mutations are stratified into CpG and non-CpG contexts.

To understand the specific base preferences driving these *RE* statistics, we examined the directional shifts in nucleotide frequencies at flanking sites between the singletons and their location-matched control sites. For the *A→G* mutation sub-type, which exhibited the highest *RE* statistic at the +1 position, we observed a stark divergence in nucleotide distributions highlighted by a difference in the proportion of thymine flanking singletons (*p*_*T*_ = 0.427) compared to the genomic background (*p*_*T*_ = 0.265). This 1.6-fold enrichment of thymine at the +1 position represents an excess of approximately 2.33 million *A→G* variants associated with a +1 T-neighbor relative to the null expectation. Conversely, the smallest *RE* within the ± 10 bp window—found at the − 7 position for the *C→T* subtype corresponds to a much subtler shift (singleton *p*_*C*_ = 0.232 vs. control *p*_*C*_ = 0.230). Despite this representing only a 1% relative change in frequency, the high power of our dataset identifies this as a significant shift corresponding to 29,056 ‘excess’ singletons. These examples illustrate that even minute shifts in local sequence context can correspond to substantial variations in mutational load across the genome.

While these raw frequency shifts provide concrete examples of local context effects, systematically comparing directional biases across all flanking positions requires evaluating the nucleotide-level residuals from our log-linear models. Because of our 5-to-1 ratio of controls to singletons, this estimated shared distribution is dominated by the background controls, with the resulting residuals revealing where the mutation processes diverge from background genomic contexts. Among *A*? *N* mutations, we observed a shared contextual signature for *A*? *G* and *A*? *T* substitutions, both characterized by a marked enrichment of T and a depletion of A and G at the +1 position. This pattern points to a common mutational pressure for these classes that is entirely absent in *A*? *C* transversions, where the +1 signal was instead driven by an enrichment of A alongside a depletion of T and C.

Distinct patterns also emerged among the non-CpG *C→N* subtypes. Unlike the *A*? *N* group, no single flanking position uniformly dominated the *RE* signal across the three *C→N* subtypes, and the underlying nucleotide residuals were highly variable. For instance, at the +1 position, *C→A* mutations were characterized by an enrichment of C and a depletion of T, whereas the +1 signal for *C→G* was primarily driven by a depletion of A and an enrichment of T. In contrast, the strongest signals observed in the CpG subtypes were all observed at -1; however, the driver of the underlying signal was distinct across the three subtypes, with an enrichment of G characterizing the signal for the *CpG→ApG* subtype, a depletion of G for the *CpG* ?*GpG* subtype, and a depletion of T for the *CpG→ TpG* subtype. Altogether, these contrasting residual patterns underscore substantial heterogeneity in sequence-context sensitivity across different base substitutions at Cytosine sites.

Although relative entropy generally decays as distance from the focal site increases, all deviance statistics within the initial ± 10 bp window remained highly significant across all mutation subtypes (*p <* 7 × 10^−6^). To determine the spatial limit of this influence, we extended our analysis to evaluate marginal effects at positions up to 1, 000 bp from the focal position. To safeguard against false positives, we established an empirical null distribution by comparing nucleotide frequencies between independent, non-overlapping sets of matched control sites. As illustrated by the *A→G* subtype (Figure 2), the most substantial *RE* values are heavily concentrated near the focal site, however, statistically significant deviations persist well beyond the traditional ± 10 bp boundary.

**Fig. 2:**
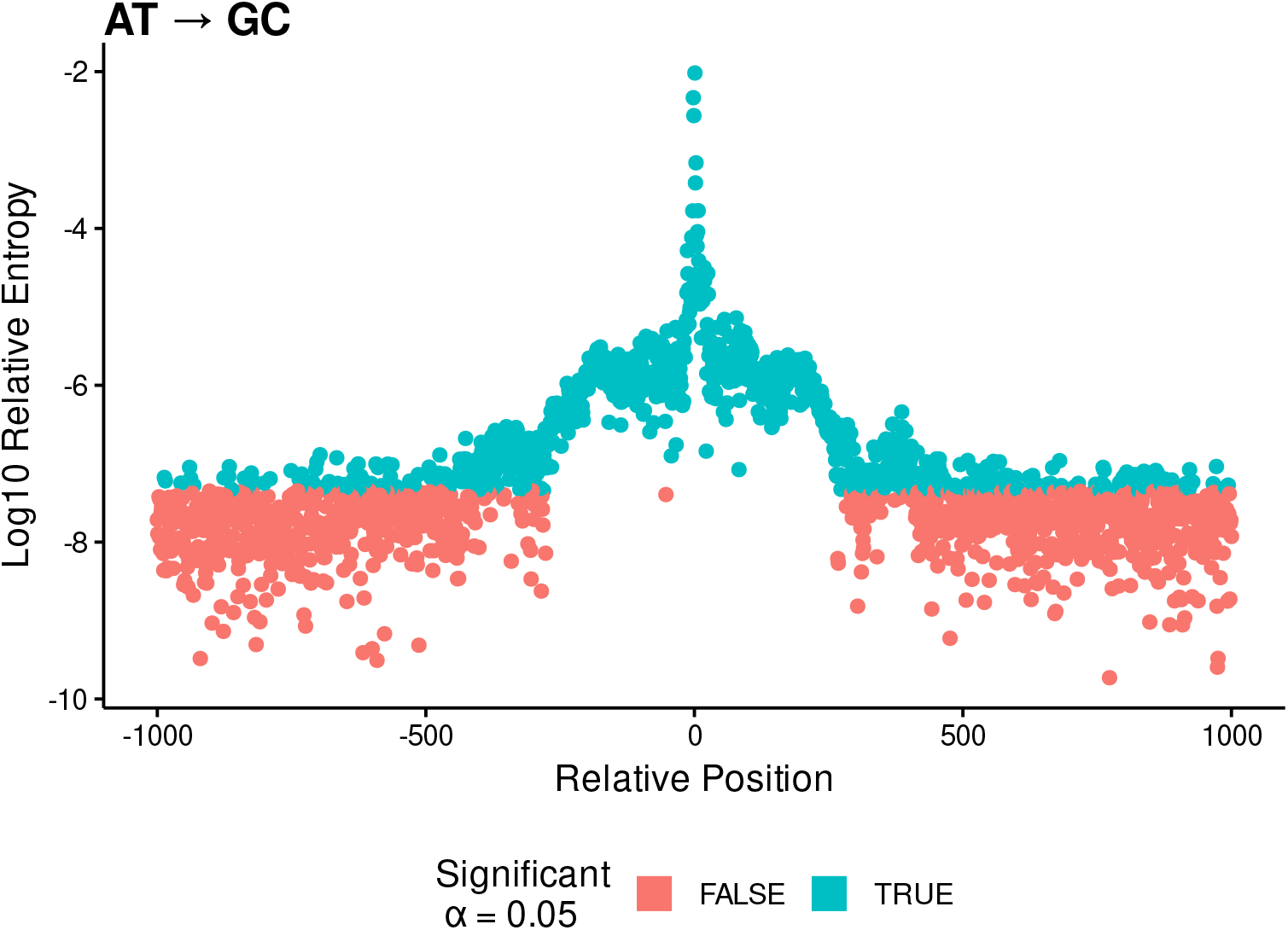
Long-range spatial decay of single-position flanking influence for the *A→G* mutation subtype. Relative entropy (*RE*) statistics are plotted for individual flanking positions up to 1, 000 bp from the focal mutation site.

To determine the spatial boundary of these long-range context effects, we used a symmetric sliding window to identify the coordinate range within which 95% of flanking positions maintained statistically significant deviance (*α* = 0.05). For the *A→ G* subtype, this threshold reached ±300 bp from the focal position. We observed similar ranges across the other non-CpG subtypes, with this “influence horizon” consistently falling between ±295 and ±335 bp (Supplementary Table 3). The boundaries derived via this sliding-window approach align with the spatial coordinates at which the observed *RE* statistics converge with our empirical null distributions, providing a data-driven definition of the outer limit of single-position flanking influence.

### 3.2 Pairwise interactions among flanking nucleotides

Having established that individual nucleotides appear to influence mutation rates across several hundred base pairs, we next evaluated whether combinations of nucleotides at pairs of flanking positions affect the mutation process beyond what can be predicted based on their marginal effects. For all subtypes, interaction strength (quantified by *RE*) was inversely related to the distance between the two positions and their distances from the focal site. All but one subtype exhibited its peak two-way interaction at pairs involving at least one position immediately adjacent to the focal site (± 1 bp), and for seven of the nine subtypes the maximum two-way signal was contained within a ± 2 bp radius of the focal site (Figure 3). The CpG ?TpG sub-type was the lone exception to this pattern, with its peak two-position effect observed at the (+3, +4) positions.

**Fig. 3:**
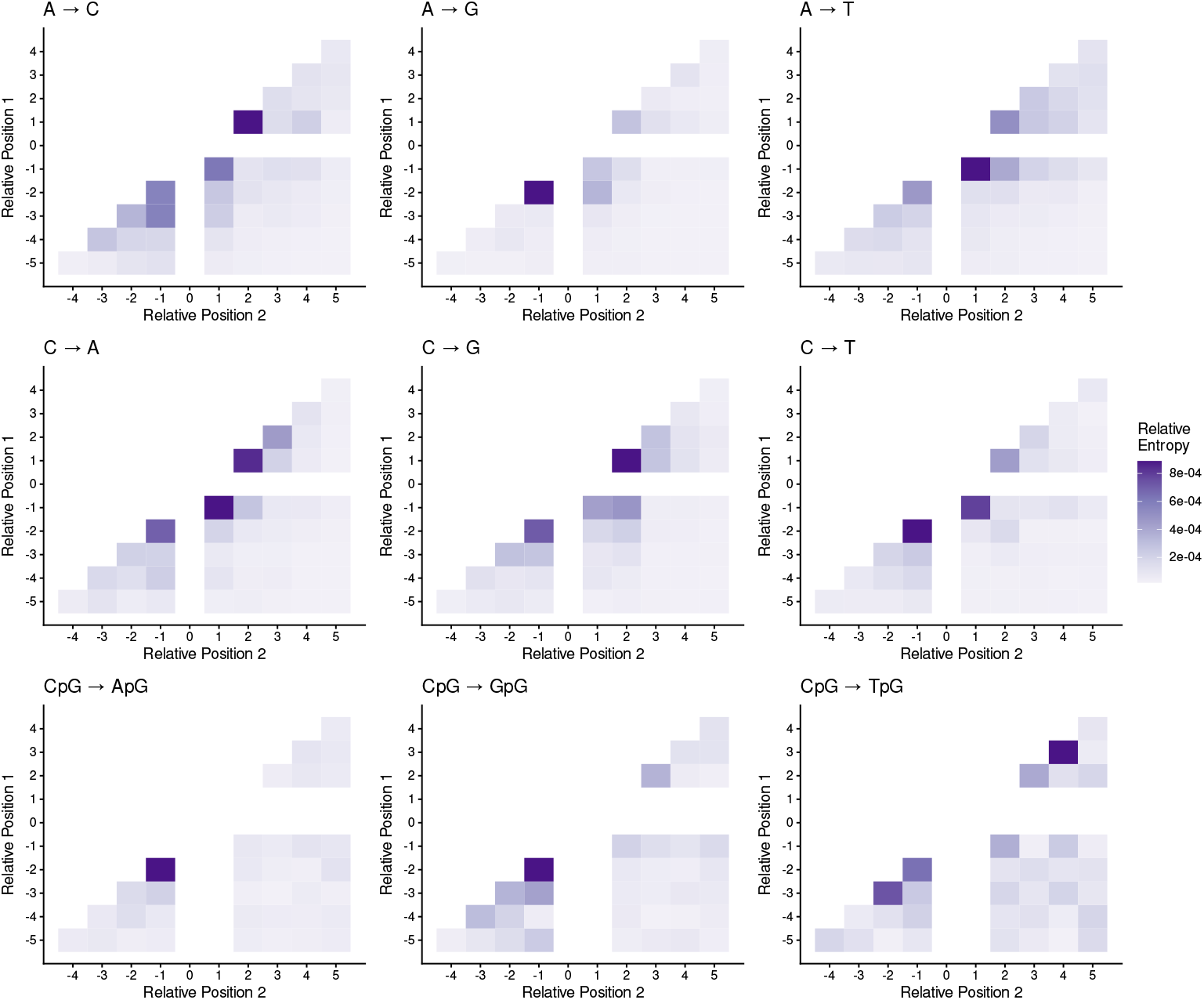
Two-position RE values observed at pairs of flanking positions within a ± 5*bp* window for each mutation subtype.

**Fig. 4:**
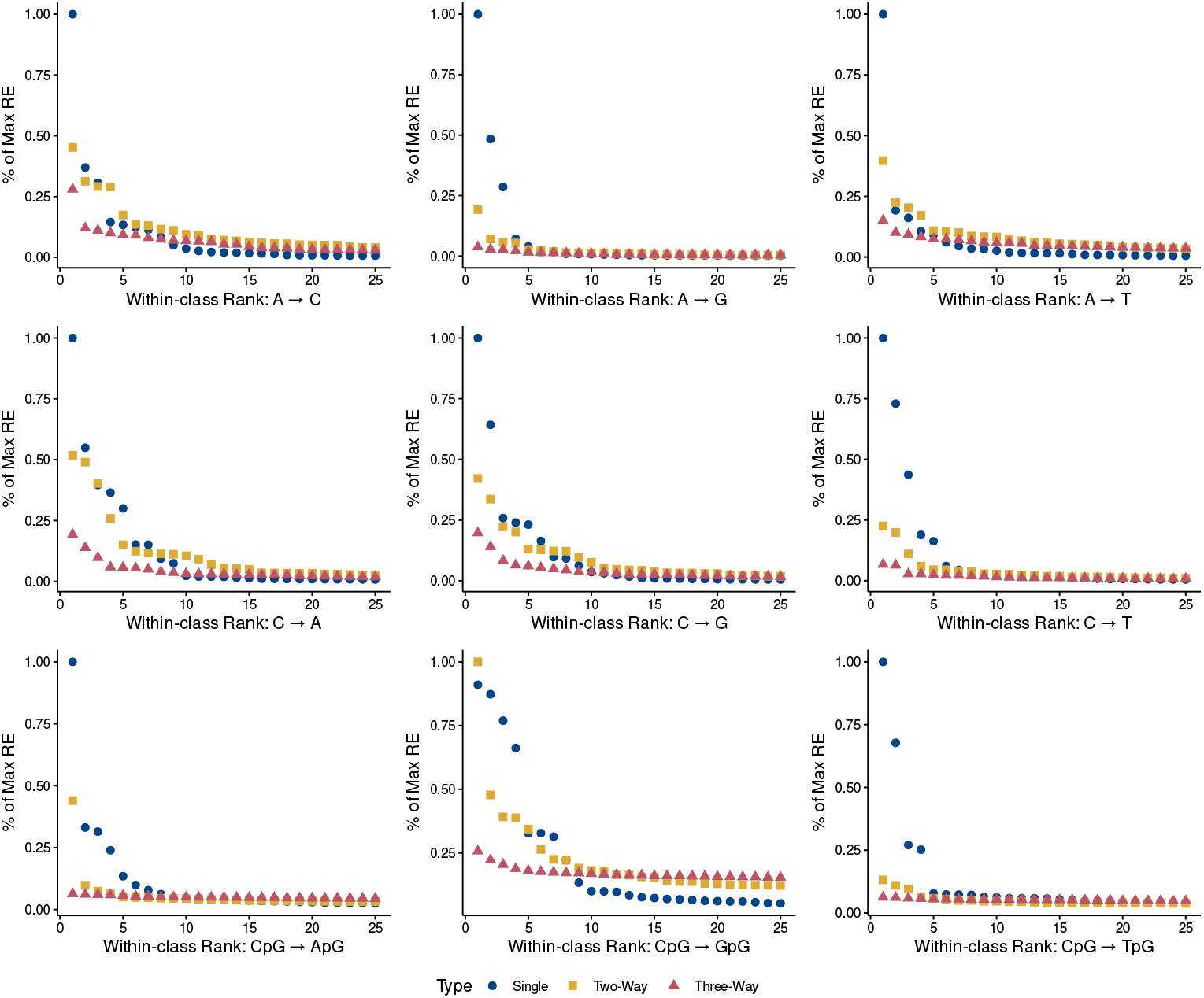
Relative entropy decay across single-, two-, and three-position model classes. Within-class ranks (x-axis; ranks 1–20) for single-position, two-position, and three-position models plotted against the normalized relative entropy percentage (% max *RE*; y-axis) relative to the top-ranked effect in each mutation subtype. In three of nine subtypes (*A→C, A→T*, and CpG→A), the top-ranked second-order interaction constitutes the second strongest signal overall (∼ 50% of maximum *RE*), while in CpG→G, a second-order term defines the 100% maximum *RE*. In the remaining five subtypes, one to three single-position main effects exceed the top pairwise interaction in magnitude. Across all subtypes, third-order interactions drop off sharply, rarely exceeding 25% of maximum *RE*.

Just as marginal *RE* statistics can be decomposed into nucleotide-level contributions, two-way interaction terms can be partitioned into specific dinucleotide pairs to evaluate non-additive directional effects. We illustrate this using the *A→G* subtype, where the strongest two-way interaction occurred at the (− 2, −1) positions. In our single-position analyses for this subtype, the − 2 position was primarily characterized by an enrichment of C in singletons, while the − 1 position was driven by a depletion of G. After explicitly accounting for these first-order main effects, we identified a significant non-additive effect characterized by the co-occurrence of C and A at (− 2, − 1) being enriched in singletons, whereas the C and C pairing is depleted. This indicates that the presence of a − 1 adenine further amplifies the mutability already associated with a − 2 cytosine, whereas a − 1 cytosine appears to mitigate it. Such dependencies reveal that the influences of flanking nucleotides on mutation rates are themselves often dependent on local sequence context.

### 3.3 Assessing the relative strength of higher order effects

To determine the relative contributions of context dependencies across model complexities, we compared the magnitude and spatial distribution of first-, second-, and third-order effects across all mutation subtypes. Overall, we observed a general hierarchical decay in signal magnitude as interaction complexity increased. With the exception of the CpG→*G* subtype—where a two-way interaction constituted the single strongest signal overall—the peak *RE* statistic for every class was driven by a single-position marginal effect. When comparing second-order interactions against marginal effects, subtypes fell into distinct regimes: in three classes (*A*? *C, A*? *T*, and CpG → ApG), the peak pairwise interaction represented the second strongest overall signal (reaching ∼ 50% of the lead main effect), whereas in five classes, the top pairwise signal was outranked by multiple marginal effects. Finally, third-order dependencies demonstrated extreme spatial localization. While marginal and pairwise signals extended further into flanking sequence, the peak three-way interactions for non-CpG subtypes were strictly restricted to contiguous triplets immediately adjacent to or straddling the focal site, such as {−2, −1, +1} for *A→G* and non-CpG *C→T*, and{−3, −2, −1}for *C→G* (Suppl. Figure 7).

To compare the relative contributions of first-, second-, and third-order effects across subtypes, we restricted our attention to single sites, pairs, and trios whose *RE* statistics were at least 5% of the maximum *RE* statistic observed within that sub-type [8]. Under this threshold, the total number of surviving terms and their spatial scope varied dramatically across mutation subtypes. Among non-CpG subtypes, context dependencies operated within tight, highly localized boundaries. Subtypes such as *A→C* and *A→T* retained substantial higher-order structure, maintaining up to 22 pairwise and 14 three-way interactions within compact windows ([−8, +4] bp and [−4, +5] bp, respectively). Conversely, *A→G* and non-CpG *C→T* collapsed almost entirely to single-position dominance: *A→G* retained zero three-way terms within a narrow [−2, +3] bp span, while *C→T* maintained just 4 pairwise and 2 three-way terms within [− 3, +4] bp. *C→A* and *C→G* exhibited intermediate profiles, maintaining 6 to 7 three-way terms strictly bounded within [− 5, +5] bp. In contrast, CpG subtypes displayed far broader spatial windows. In CpG→ApG, surviving effects expanded across a [− 10, +10] bp window, retaining 10 three-way terms. CpG→*T* maintained 19 three-way terms spanning an even wider [− 28, +35] bp region, although all three-way effects were no larger than 6.3% of peak marginal *RE*. CpG→GpG exhibited a unique architecture due to its low baseline maximum *RE*, with 317 pair-wise and 918 three-way interactions meeting the 5% relative threshold, spreading across an expansive [−49, +55] bp window.

### 3.4 Patterns of local sequence context influence on mutation are consistent across 1kGP populations

We next assessed the consistency of sequence-dependent mutability across the distinct populations represented in the 1kGP dataset. Across all nine mutation subtypes, marginal *RE* values within ± 20 bp of the focal position were nearly identical between populations (Pearson *ρ* range: 0.959–1.00, mean: 0.996, Figure 8). This high degree of concordance persisted even when the window was expanded to ± 1, 000 bp (mean *ρ* = 0.995). Similarly, the second-order interactions demonstrated high stability across populations within the ± 20 bp window (Pearson *ρ* mean: 0.98; Spearman *ρ* mean: 0.86).

The spatial distribution of these signals was also highly conserved, as the ranking of the most influential flanking positions was consistent across populations for all but the *CpG→GpG* subtype, where the peak influence was located at the +2 position for the AFR population but at the − 1 position for all other groups (Figure 9). Furthermore, when decomposing these statistics into specific nucleotide-level residuals, we observed in every subtype at the most influential sites the nucleotides driving the primary enrichments and depletions were shared across all populations.

## 4 Discussion

In this study, we leveraged a high-coverage dataset of autosomal singleton variants from the 1000 Genomes Project (1kGP) within an information-theoretic framework to construct a comprehensive map of germline sequence-dependent mutability. By focusing on evolutionarily recent singletons, our analysis effectively removed post-mutational confounders such as natural selection and biased gene conversion. Beyond confirming the baseline dominance of immediate single-position effects [8], our log-linear decomposition yielded several insights into the architecture of germline context dependencies. We significantly expanded the genomic window of analysis to demonstrate that individual flanking positions exert a subtle but statistically robust influence extending up to several hundred base pairs. Higher-order interactions were also evaluated, and while influence generally decayed as interaction complexity increased, second- and third-order effects proved influential, further differentiating patterns observed across mutation subtypes. Finally, we demonstrated high concordance in both single position and higher-order interaction effects across all 1kGP super-populations, confirming that these patterns of local sequence context influence on germline mutation rates reflect conserved features of the human germline mutational machinery.

Prior efforts to model local germline mutation rates leveraged the distribution of non-coding variants to describe the biological footprints of germline mutagenesis [7, 8]. While [9] subsequently demonstrated that singletons from a large sample of unrelated subjects better reflect these signatures by minimizing confounding from other forces such as natural selection and biased GC-conversion, their strategy of fitting separate 7-mer models with genomic features left local context effects partially entangled with broader genomic features that are often difficult to comprehensively measure. Here, by applying a log-linear model comparing the distributions of nucleotides flanking singletons to a genomic location-matched control distribution [8], we cleanly isolate sequence context from regional genomic confounders. Furthermore, our decomposition of single-position main effects versus higher-order interactions provides empirical justification for recent extended-context models such as *Roulette* [10], which incorporated local sequence context in a broader window in their germline mutation rate model by adding only main effect terms for flanking positions outside of the interior pentamer. By demonstrating that the influence of interactions decreases with both order and distance from the focal site, our results establish an empirical basis for extended flanking contexts via single-position effects in a manner that avoids exponential parameter growth.

While conventional mutational models typically restrict sequence context to immediate flanking nucleotides within 3-to 7-mer windows [7, 8], individual flanking bases exert a subtle but statistically significant influence extending across several hundred base pairs in ranges that vary by mutation subtype. A central finding of our expanded spatial analysis is the marked contrast between the broad reach of single-position marginal effects and the localized footprint of higher-order interactions. Across all effect orders, context influence generally decays both as interaction complexity increases and as spatial distance from the focal site expands. This spatial and hierarchical decay provides an empirical justification for extended-context frameworks, such as *Roulette* [10], which incorporate flanking positions beyond a central 5-mer using only main-effect terms. By isolating sequence context from regional genomic confounders through location-matched controls [9], our results formally demonstrate that expanding contextual models via single-position main effects captures the broad additive sequence landscape, likely reflecting regional structural properties of the DNA duplex, without incurring the exponential parameter growth required by full higher-order parameterizations.

Beyond improving our understanding of baseline germline mutability, these findings have immediate practical implications for how sequence context is parameterized in predictive and signature-based mutation models. For example, the COSMIC project [13] defines mutational signatures by deconvolving observed variants across centered 3-mer mutation spectra, with key single-base substitution signatures (such as SBS1 and SBS5) mirroring patterns observed in germline mutagenesis [14]. While 3-mer representations are computationally efficient, our evaluation reveals a fundamental limitation in how these motifs are conventionally constructed. Existing frameworks implicitly assume that local epistatic interactions are universally anchored at the immediate (− 1, +1) positions. However, we demonstrate that peak pairwise dependencies are subtype-specific and frequently off-center relative to the focal site—exemplified by *A*? *G*, where the primary interaction occurs between the upstream (− 2, − 1) pair. Consequently, the computational tractability of a 3-mer representation could be preserved while gaining substantial explanatory power simply by adopting flexible, subtype-specific 3-mer windows rather than a rigid symmetric template. Furthermore, by restricting context strictly to immediate neighbors, standard 3-mer models miss the cumulative long-range marginal influences that span tens to hundreds of base pairs. Incorporating both asymmetric local interaction windows and extended flanking main effects will be essential for refining the next generation of germline and somatic mutation models.

While our framework establishes a clear quantification of sequence-dependent mutability, local nucleotide context operates within a broader, hierarchical genomic architecture in moderating germline mutagenesis. By utilizing a location-matched control sampling strategy [8], our framework explicitly controls for large-scale genomic covariates, such as regional GC content, replication timing, histone modifications, and recombination rates, allowing us to isolate the mutational footprint of germline mutagenesis from regional variation. In reality, these regional features can modulate local repair efficiency and polymerase fidelity [5, 15]. For example, late-replicating regions and heterochromatic domains frequently exhibit elevated baseline substitution rates. A key question moving forward is whether the long-range marginal effects observed in our model act independently of this regional substrate, or if specific flanking motifs dynamically amplify mutability in distinct epigenomic environments. Integrating our multi-position contextual parameters with fine-mapped epigenomic and replicationtiming tracks will be a crucial next step toward building fully unified, multi-scale models of germline mutability.

Our cross-population analyses demonstrate remarkable concordance in both marginal effects (Pearson *ρ* ≥ 0.959) and higher-order interactions (mean Pearson *ρ* = 0.98) across all 1kGP super-populations, pointing towards a shared architecture of context-dependent mutability across modern human populations. In recent human history, transient shifts in specific mutational signatures have been documented, most notably a localized burst in *TCC→TTC* mutations in ancestral West Eurasian populations [16, 17]. However, our reliance on rare autosomal singletons isolates recent germline events across global populations, hinting that such population-specific pulses represent localized deviations from an otherwise stable mutational background. The consistency observed across all nine substitution subtypes suggests that while transient modifiers or environmental factors may periodically alter the rate of specific local contexts, the underlying human germline mutational machinery is quite similar across modern human populations.

By leveraging the distribution of rare autosomal singletons across a global population sample, our work provides a detailed picture of how local sequence context shapes germline mutagenesis. Our findings suggest a conceptual shift of moving away from treating local sequence context as a rigid, symmetric *k*-mer motif, and toward an architecture that balances localized, non-linear epistatic interactions focal to the site of mutation with additive, long-range flanking influences. Decoupling these localized non-additive interactions from extended main effects offers a principled solution to a long-standing model-design trade-off, enabling high-resolution mutation models that maximize predictive power without incurring exponential parameter growth. As large-scale sequencing cohorts continue to expand, incorporating these flexible context architectures will be vital for disentangling baseline mutational processes from natural selection, refining clinical disease-variant prioritization, and building fully unified models of genome evolution.

## 5 Supplementary Tables and Figures

### 5.1 Tables

**Table 1:** Deviance statistics, relative entropy, and p-values for all flanking positions within 10 bases of the focal position across all mutation subtypes. C* denotes subtypes where we don’t distinguish between CpG and non-CpG mutations, while C denotes subtypes where we’ve excluded CpG sites.

| Subtype | Offset | Deviance | Relative Entropy | P value |
| --- | --- | --- | --- | --- |
| A → C | -10 | 9.845248e+02 | 0.0000189 | 0.0000000 |
| A → C | -9 | 1.927368e+03 | 0.0000371 | 0.0000000 |
| A → C | -8 | 8.545480e+03 | 0.0001644 | 0.0000000 |
| A → C | -7 | 2.686660e+03 | 0.0000517 | 0.0000000 |
| A → C | -6 | 2.351322e+02 | 0.0000045 | 0.0000000 |
| A → C | -5 | 1.735946e+03 | 0.0000334 | 0.0000000 |
| A → C | -4 | 1.249387e+04 | 0.0002403 | 0.0000000 |
| A → C | -3 | 3.772837e+04 | 0.0007257 | 0.0000000 |
| A → C | -2 | 5.035458e+03 | 0.0000969 | 0.0000000 |
| A → C | -1 | 1.020876e+05 | 0.0019635 | 0.0000000 |
| A → C | 1 | 1.365441e+04 | 0.0002626 | 0.0000000 |
| A → C | 2 | 3.128926e+04 | 0.0006018 | 0.0000000 |
| A → C | 3 | 1.481820e+04 | 0.0002850 | 0.0000000 |
| A → C | 4 | 1.161563e+04 | 0.0002234 | 0.0000000 |
| A → C | 5 | 3.650904e+03 | 0.0000702 | 0.0000000 |
| A → C | 6 | 8.739774e+02 | 0.0000168 | 0.0000000 |
| A → C | 7 | 8.481566e+02 | 0.0000163 | 0.0000000 |
| A → C | 8 | 1.466236e+03 | 0.0000282 | 0.0000000 |
| A → C | 9 | 2.050471e+03 | 0.0000394 | 0.0000000 |
| A → C | 10 | 3.133489e+02 | 0.0000060 | 0.0000000 |
| A → G | -10 | 1.039506e+03 | 0.0000060 | 0.0000000 |
| A → G | -9 | 1.478470e+03 | 0.0000086 | 0.0000000 |
| A → G | -8 | 1.698000e+03 | 0.0000098 | 0.0000000 |
| A → G | -7 | 2.730530e+03 | 0.0000158 | 0.0000000 |
| A → G | -6 | 2.069945e+03 | 0.0000120 | 0.0000000 |
| A → G | -5 | 1.327931e+04 | 0.0000768 | 0.0000000 |
| A → G | -4 | 3.059636e+03 | 0.0000177 | 0.0000000 |
| A → G | -3 | 2.898218e+04 | 0.0001677 | 0.0000000 |
| A → G | -2 | 8.030187e+05 | 0.0046464 | 0.0000000 |
| A → G | -1 | 4.745049e+05 | 0.0027455 | 0.0000000 |
| A → G | 1 | 1.659567e+06 | 0.0096025 | 0.0000000 |
| A → G | 2 | 6.586906e+04 | 0.0003811 | 0.0000000 |
| A → G | 3 | 1.186398e+05 | 0.0006865 | 0.0000000 |
| A → G | 4 | 1.418849e+04 | 0.0000821 | 0.0000000 |
| A → G | 5 | 1.025300e+04 | 0.0000593 | 0.0000000 |
| A → G | 6 | 1.570967e+04 | 0.0000909 | 0.0000000 |
| A → G | 7 | 2.911467e+04 | 0.0001685 | 0.0000000 |
| A → G | 8 | 6.725429e+03 | 0.0000389 | 0.0000000 |
| A → G | 9 | 2.947974e+03 | 0.0000171 | 0.0000000 |
| A → G | 10 | 1.876655e+03 | 0.0000109 | 0.0000000 |
| A → T | -10 | 1.262900e+03 | 0.0000266 | 0.0000000 |
| A → T | -9 | 1.182325e+03 | 0.0000249 | 0.0000000 |
| A → T | -8 | 1.901583e+03 | 0.0000400 | 0.0000000 |
| A → T | -7 | 2.582574e+03 | 0.0000543 | 0.0000000 |
| A → T | -6 | 3.632761e+03 | 0.0000764 | 0.0000000 |
| A → T | -5 | 7.331928e+03 | 0.0001541 | 0.0000000 |
| A → T | -4 | 1.285975e+04 | 0.0002704 | 0.0000000 |
| A → T | -3 | 9.518656e+03 | 0.0002001 | 0.0000000 |
| A → T | -2 | 1.954573e+04 | 0.0004109 | 0.0000000 |
| A → T | -1 | 4.077034e+04 | 0.0008571 | 0.0000000 |
| A → T | 1 | 2.120337e+05 | 0.0044577 | 0.0000000 |
| A → T | 2 | 3.420457e+04 | 0.0007191 | 0.0000000 |
| A → T | 3 | 2.243154e+04 | 0.0004716 | 0.0000000 |
| A → T | 4 | 5.482603e+03 | 0.0001153 | 0.0000000 |
| A → T | 5 | 6.879559e+03 | 0.0001446 | 0.0000000 |
| A → T | 6 | 3.266340e+03 | 0.0000687 | 0.0000000 |
| A → T | 7 | 4.123214e+03 | 0.0000867 | 0.0000000 |
| A → T | 8 | 3.361220e+03 | 0.0000707 | 0.0000000 |
| A → T | 9 | 1.556080e+03 | 0.0000327 | 0.0000000 |
| A → T | 10 | 1.801168e+03 | 0.0000379 | 0.0000000 |
| C → T | -10 | 8.360217e+02 | 0.0000053 | 0.0000000 |
| C → T | -9 | 2.589075e+03 | 0.0000164 | 0.0000000 |
| C → T | -8 | 1.659288e+03 | 0.0000105 | 0.0000000 |
| C → T | -7 | 4.305808e+02 | 0.0000027 | 0.0000000 |
| C → T | -6 | 1.963832e+03 | 0.0000124 | 0.0000000 |
| C → T | -5 | 1.073939e+04 | 0.0000680 | 0.0000000 |
| C → T | -4 | 5.147413e+03 | 0.0000326 | 0.0000000 |
| C → T | -3 | 3.902650e+03 | 0.0000247 | 0.0000000 |
| C → T | -2 | 3.955112e+04 | 0.0002504 | 0.0000000 |
| C → T | -1 | 2.432812e+05 | 0.0015401 | 0.0000000 |
| C → T | 1 | 1.062135e+05 | 0.0006724 | 0.0000000 |
| C → T | 2 | 1.774909e+05 | 0.0011236 | 0.0000000 |
| C → T | 3 | 4.596741e+04 | 0.0002910 | 0.0000000 |
| C → T | 4 | 1.470394e+04 | 0.0000931 | 0.0000000 |
| C → T | 5 | 5.663166e+03 | 0.0000359 | 0.0000000 |
| C → T | 6 | 3.393586e+03 | 0.0000215 | 0.0000000 |
| C → T | 7 | 5.006380e+03 | 0.0000317 | 0.0000000 |
| C → T | 8 | 4.018413e+03 | 0.0000254 | 0.0000000 |
| C → T | 9 | 4.881638e+03 | 0.0000309 | 0.0000000 |
| C → T | 10 | 6.964804e+03 | 0.0000441 | 0.0000000 |
| C → G | -10 | 3.040730e+02 | 0.0000056 | 0.0000000 |
| C → G | -9 | 1.234094e+03 | 0.0000229 | 0.0000000 |
| C → G | -8 | 1.098777e+03 | 0.0000204 | 0.0000000 |
| C → G | -7 | 9.464860e+02 | 0.0000175 | 0.0000000 |
| $C \rightarrow G$ | -6 | 3.373733e+03 | 0.0000625 | 0.0000000 |
| $C \rightarrow G$ | -5 | 9.206343e+03 | 0.0001705 | 0.0000000 |
| $C \rightarrow G$ | -4 | 1.430293e+04 | 0.0002649 | 0.0000000 |
| $C \rightarrow G$ | -3 | 2.394901e+04 | 0.0004436 | 0.0000000 |
| $C \rightarrow G$ | -2 | 1.461141e+05 | 0.0027065 | 0.0000000 |
| $C \rightarrow G$ | -1 | 3.771763e+04 | 0.0006987 | 0.0000000 |
| $C \rightarrow G$ | 1 | 9.389886e+04 | 0.0017393 | 0.0000000 |
| $C \rightarrow G$ | 2 | 3.378497e+04 | 0.0006258 | 0.0000000 |
| $C \rightarrow G$ | 3 | 3.500464e+04 | 0.0006484 | 0.0000000 |
| $C \rightarrow G$ | 4 | 4.526636e+03 | 0.0000838 | 0.0000000 |
| $C \rightarrow G$ | 5 | 1.343346e+04 | 0.0002488 | 0.0000000 |
| $C \rightarrow G$ | 6 | 5.274599e+03 | 0.0000977 | 0.0000000 |
| $C \rightarrow G$ | 7 | 2.531277e+03 | 0.0000469 | 0.0000000 |
| $C \rightarrow G$ | 8 | 6.503669e+02 | 0.0000120 | 0.0000000 |
| $C \rightarrow G$ | 9 | 1.384108e+03 | 0.0000256 | 0.0000000 |
| $C \rightarrow G$ | 10 | 1.633973e+02 | 0.0000030 | 0.0000000 |
| $C \rightarrow A$ | -10 | 1.670548e+03 | 0.0000279 | 0.0000000 |
| $C \rightarrow A$ | -9 | 2.484537e+03 | 0.0000416 | 0.0000000 |
| $C \rightarrow A$ | -8 | 4.407485e+03 | 0.0000737 | 0.0000000 |
| $C \rightarrow A$ | -7 | 1.646602e+03 | 0.0000275 | 0.0000000 |
| $C \rightarrow A$ | -6 | 1.978236e+03 | 0.0000331 | 0.0000000 |
| $C \rightarrow A$ | -5 | 2.078893e+04 | 0.0003477 | 0.0000000 |
| $C \rightarrow A$ | -4 | 3.346774e+04 | 0.0005598 | 0.0000000 |
| $C \rightarrow A$ | -3 | 6.659887e+04 | 0.0011139 | 0.0000000 |
| $C \rightarrow A$ | -2 | 1.219258e+05 | 0.0020393 | 0.0000000 |
| $C \rightarrow A$ | -1 | 8.799978e+04 | 0.0014719 | 0.0000000 |
| $C \rightarrow A$ | 1 | 2.220352e+05 | 0.0037137 | 0.0000000 |
| $C \rightarrow A$ | 2 | 8.099892e+04 | 0.0013548 | 0.0000000 |
| $C \rightarrow A$ | 3 | 3.362171e+04 | 0.0005624 | 0.0000000 |
| $C \rightarrow A$ | 4 | 1.647182e+04 | 0.0002755 | 0.0000000 |
| $C \rightarrow A$ | 5 | 4.122994e+03 | 0.0000690 | 0.0000000 |
| $C \rightarrow A$ | 6 | 6.719081e+02 | 0.0000112 | 0.0000000 |
| $C \rightarrow A$ | 7 | 6.211088e+02 | 0.0000104 | 0.0000000 |
| $C \rightarrow A$ | 8 | 3.164840e+02 | 0.0000053 | 0.0000000 |
| $C \rightarrow A$ | 9 | 9.935191e+02 | 0.0000166 | 0.0000000 |
| $C \rightarrow A$ | 10 | 2.193343e+03 | 0.0000367 | 0.0000000 |
| $C^* \rightarrow T$ | -10 | 1.916293e+03 | 0.0000081 | 0.0000000 |
| $C^* \rightarrow T$ | -9 | 1.385250e+04 | 0.0000584 | 0.0000000 |
| $C^* \rightarrow T$ | -8 | 2.095243e+02 | 0.0000009 | 0.0000000 |
| $C^* \rightarrow T$ | -7 | 1.471577e+04 | 0.0000620 | 0.0000000 |
| $C^* \rightarrow T$ | -6 | 3.670557e+03 | 0.0000155 | 0.0000000 |
| $C^* \rightarrow T$ | -5 | 9.921012e+03 | 0.0000418 | 0.0000000 |
| $C^* \rightarrow T$ | -4 | 3.662257e+03 | 0.0000154 | 0.0000000 |
| $C^* \rightarrow T$ | -3 | 1.331525e+04 | 0.0000561 | 0.0000000 |
| $C^* \rightarrow T$ | -2 | 8.355645e+04 | 0.0003521 | 0.0000000 |
| $C^* \rightarrow T$ | -1 | 3.962565e+05 | 0.0016699 | 0.0000000 |
| $C^* \rightarrow T$ | 1 | 1.314383e+07 | 0.0553913 | 0.0000000 |
| $C^* \rightarrow T$ | 2 | 1.011887e+05 | 0.0004264 | 0.0000000 |
| $C^* \rightarrow T$ | 3 | 5.193808e+04 | 0.0002189 | 0.0000000 |
| $C^* \rightarrow T$ | 4 | 2.093539e+04 | 0.0000882 | 0.0000000 |
| $C^* \rightarrow T$ | 5 | 1.804580e+04 | 0.0000760 | 0.0000000 |
| $C^* \rightarrow T$ | 6 | 1.554391e+04 | 0.0000655 | 0.0000000 |
| $C^* \rightarrow T$ | 7 | 8.865996e+03 | 0.0000374 | 0.0000000 |
| $C^* \rightarrow T$ | 8 | 1.262125e+04 | 0.0000532 | 0.0000000 |
| $C^* \rightarrow T$ | 9 | 4.848642e+03 | 0.0000204 | 0.0000000 |
| $C^* \rightarrow T$ | 10 | 1.378543e+04 | 0.0000581 | 0.0000000 |
| $CpG \rightarrow TpG$ | -10 | 2.822024e+03 | 0.0000356 | 0.0000000 |
| $CpG \rightarrow TpG$ | -9 | 2.658128e+03 | 0.0000335 | 0.0000000 |
| $CpG \rightarrow TpG$ | -8 | 6.067960e+03 | 0.0000765 | 0.0000000 |
| $CpG \rightarrow TpG$ | -7 | 2.169192e+03 | 0.0000273 | 0.0000000 |
| $CpG \rightarrow TpG$ | -6 | 4.825373e+03 | 0.0000608 | 0.0000000 |
| $CpG \rightarrow TpG$ | -5 | 4.822802e+03 | 0.0000608 | 0.0000000 |
| $CpG \rightarrow TpG$ | -4 | 5.180601e+03 | 0.0000653 | 0.0000000 |
| $CpG \rightarrow TpG$ | -3 | 7.260739e+02 | 0.0000092 | 0.0000000 |
| $CpG \rightarrow TpG$ | -2 | 2.223540e+04 | 0.0002803 | 0.0000000 |
| $CpG \rightarrow TpG$ | -1 | 8.209123e+04 | 0.0010349 | 0.0000000 |
| $CpG \rightarrow TpG$ | 2 | 5.562364e+04 | 0.0007012 | 0.0000000 |
| $CpG \rightarrow TpG$ | 3 | 2.069597e+04 | 0.0002609 | 0.0000000 |
| $CpG \rightarrow TpG$ | 4 | 2.040857e+03 | 0.0000257 | 0.0000000 |
| $CpG \rightarrow TpG$ | 5 | 5.131949e+03 | 0.0000647 | 0.0000000 |
| $CpG \rightarrow TpG$ | 6 | 6.013283e+03 | 0.0000758 | 0.0000000 |
| $CpG \rightarrow TpG$ | 7 | 6.426787e+03 | 0.0000810 | 0.0000000 |
| $CpG \rightarrow TpG$ | 8 | 1.926718e+03 | 0.0000243 | 0.0000000 |
| $CpG \rightarrow TpG$ | 9 | 5.905878e+03 | 0.0000745 | 0.0000000 |
| $CpG \rightarrow TpG$ | 10 | 2.688544e+03 | 0.0000339 | 0.0000000 |
| $CpG \rightarrow GpG$ | -10 | 1.864166e+01 | 0.0000036 | 0.0003242 |
| $CpG \rightarrow GpG$ | -9 | 4.091434e+02 | 0.0000784 | 0.0000000 |
| $CpG \rightarrow GpG$ | -8 | 2.979082e+02 | 0.0000571 | 0.0000000 |
| $CpG \rightarrow GpG$ | -7 | 4.086904e+02 | 0.0000783 | 0.0000000 |
| $CpG \rightarrow GpG$ | -6 | 2.834184e+02 | 0.0000543 | 0.0000000 |
| $CpG \rightarrow GpG$ | -5 | 2.160376e+03 | 0.0004140 | 0.0000000 |
| $CpG \rightarrow GpG$ | -4 | 6.863045e+02 | 0.0001315 | 0.0000000 |
| $CpG \rightarrow GpG$ | -3 | 2.255387e+03 | 0.0004322 | 0.0000000 |
| $CpG \rightarrow GpG$ | -2 | 5.293924e+03 | 0.0010144 | 0.0000000 |
| $CpG \rightarrow GpG$ | -1 | 6.262706e+03 | 0.0012000 | 0.0000000 |
| $CpG \rightarrow GpG$ | 2 | 4.553414e+03 | 0.0008725 | 0.0000000 |
| $CpG \rightarrow GpG$ | 3 | 6.007369e+03 | 0.0011511 | 0.0000000 |
| $CpG \rightarrow GpG$ | 4 | 1.526267e+03 | 0.0002925 | 0.0000000 |
| $CpG \rightarrow GpG$ | 5 | 2.254749e+03 | 0.0004321 | 0.0000000 |
| $CpG \rightarrow GpG$ | 6 | 6.826651e+02 | 0.0001308 | 0.0000000 |
| CpG → GpG | 7 | 6.690591e+02 | 0.0001282 | 0.0000000 |
| CpG → GpG | 8 | 9.185941e+02 | 0.0001760 | 0.0000000 |
| CpG → GpG | 9 | 2.534901e+02 | 0.0000486 | 0.0000000 |
| CpG → GpG | 10 | 7.932608e+01 | 0.0000152 | 0.0000000 |
| CpG → ApG | -10 | 2.689979e+01 | 0.0000041 | 0.0000062 |
| CpG → ApG | -9 | 1.027561e+03 | 0.0001547 | 0.0000000 |
| CpG → ApG | -8 | 1.272477e+03 | 0.0001916 | 0.0000000 |
| CpG → ApG | -7 | 7.700880e+02 | 0.0001159 | 0.0000000 |
| CpG → ApG | -6 | 7.616465e+02 | 0.0001147 | 0.0000000 |
| CpG → ApG | -5 | 8.848765e+03 | 0.0013323 | 0.0000000 |
| CpG → ApG | -4 | 6.731252e+03 | 0.0010135 | 0.0000000 |
| CpG → ApG | -3 | 2.783308e+03 | 0.0004191 | 0.0000000 |
| CpG → ApG | -2 | 9.311405e+03 | 0.0014019 | 0.0000000 |
| CpG → ApG | -1 | 2.810824e+04 | 0.0042319 | 0.0000000 |
| CpG → ApG | 2 | 6.554235e+02 | 0.0000987 | 0.0000000 |
| CpG → ApG | 3 | 7.067258e+02 | 0.0001064 | 0.0000000 |
| CpG → ApG | 4 | 1.751765e+03 | 0.0002637 | 0.0000000 |
| CpG → ApG | 5 | 8.441060e+02 | 0.0001271 | 0.0000000 |
| CpG → ApG | 6 | 3.783293e+03 | 0.0005696 | 0.0000000 |
| CpG → ApG | 7 | 2.197798e+03 | 0.0003309 | 0.0000000 |
| CpG → ApG | 8 | 1.086106e+03 | 0.0001635 | 0.0000000 |
| CpG → ApG | 9 | 1.216581e+03 | 0.0001832 | 0.0000000 |
| CpG → ApG | 10 | 3.735085e+02 | 0.0000562 | 0.0000000 |

**Table 2:** Counts of singletons in each mutation subtype across all 2,504 1kGP samples.

| Subtype | Singletons |
| --- | --- |
| A → C | 4332665 |
| A → G | 14402284 |
| A → T | 3963849 |
| C → A | 4982302 |
| C → G | 4498836 |
| C → T | 13163682 |
| CpG → ApG | 553495 |
| CpG → GpG | 434894 |
| CpG → TpG | 6610536 |

**Table 3:**
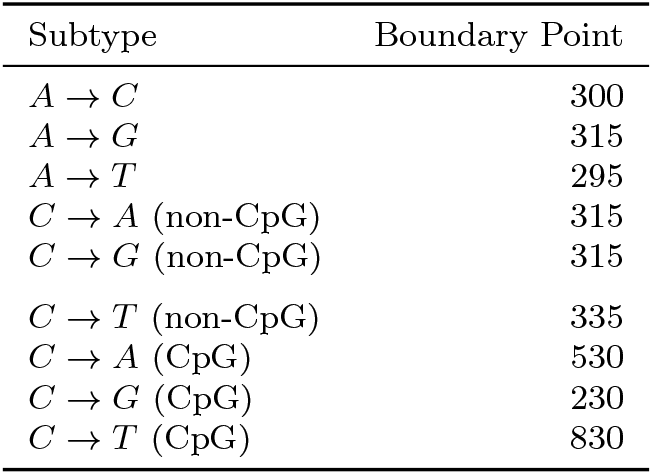
Boundary points defining the spatial extent of single-position context influences across mutation subtypes. Boundaries (in base pairs) were identified using an expanding window approach, defined as the smallest window size in which fewer than 95% of single-position test statistics reached statistical significance (*α* = 0.05). Significance at each position was evaluated against a chi-square null distribution (*χ*^2^, *df* = 3).

### 5.2 Figures

**Fig. 5:**
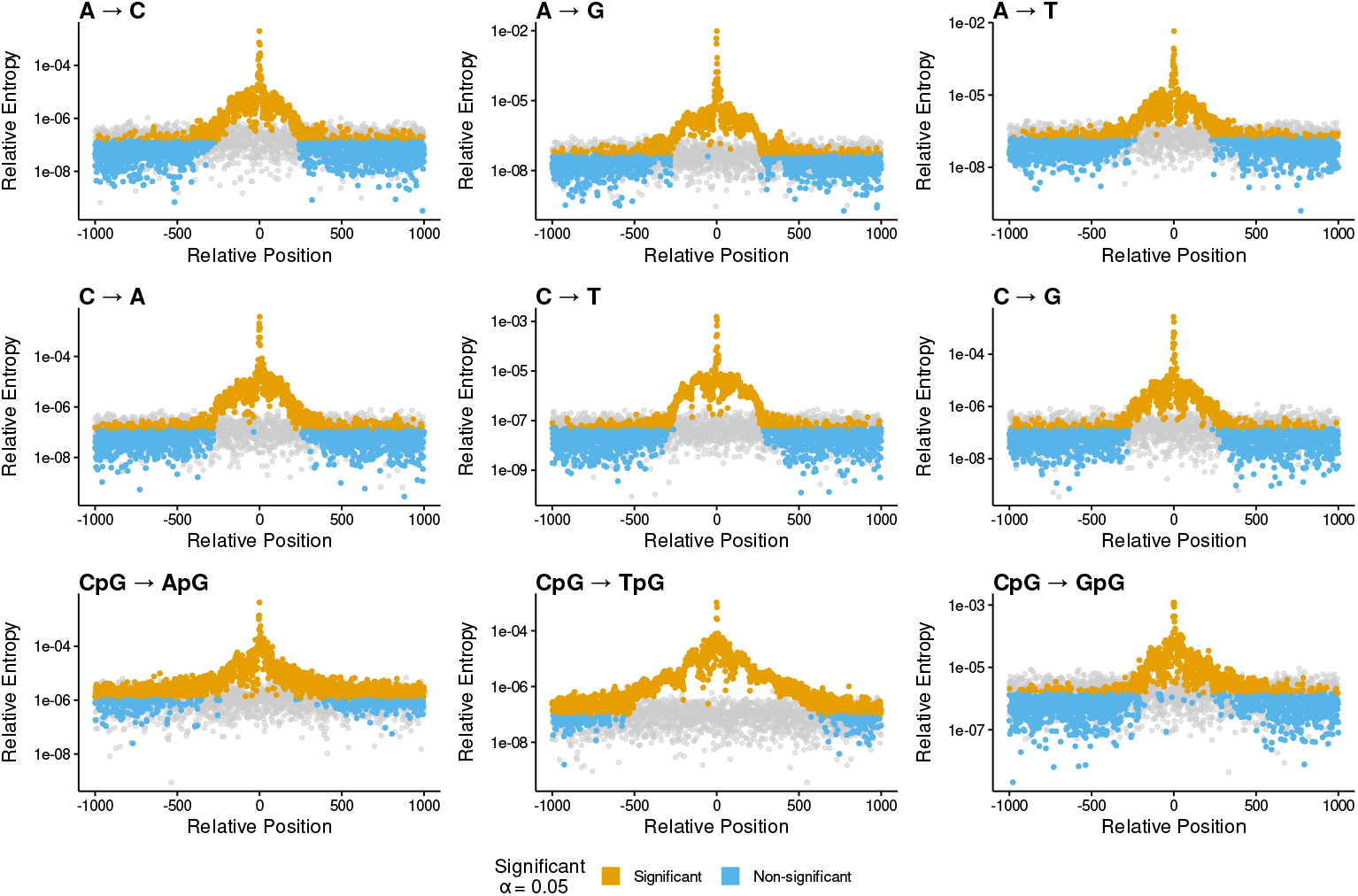
Figure S1: Genome-wide single-position mutational influence across all nine stratified mutation subtypes up to ± 1, 000 bp. Each panel displays the observed single-position relative entropy (*RE*) statistics (colored points) alongside their corresponding empirical null distributions (grey background points), generated from independent, non-overlapping sets of matched control sites. *C→N* classes are split into CpG and non-CpG contexts to control for regional methylation dynamics.

**Fig. 6:**
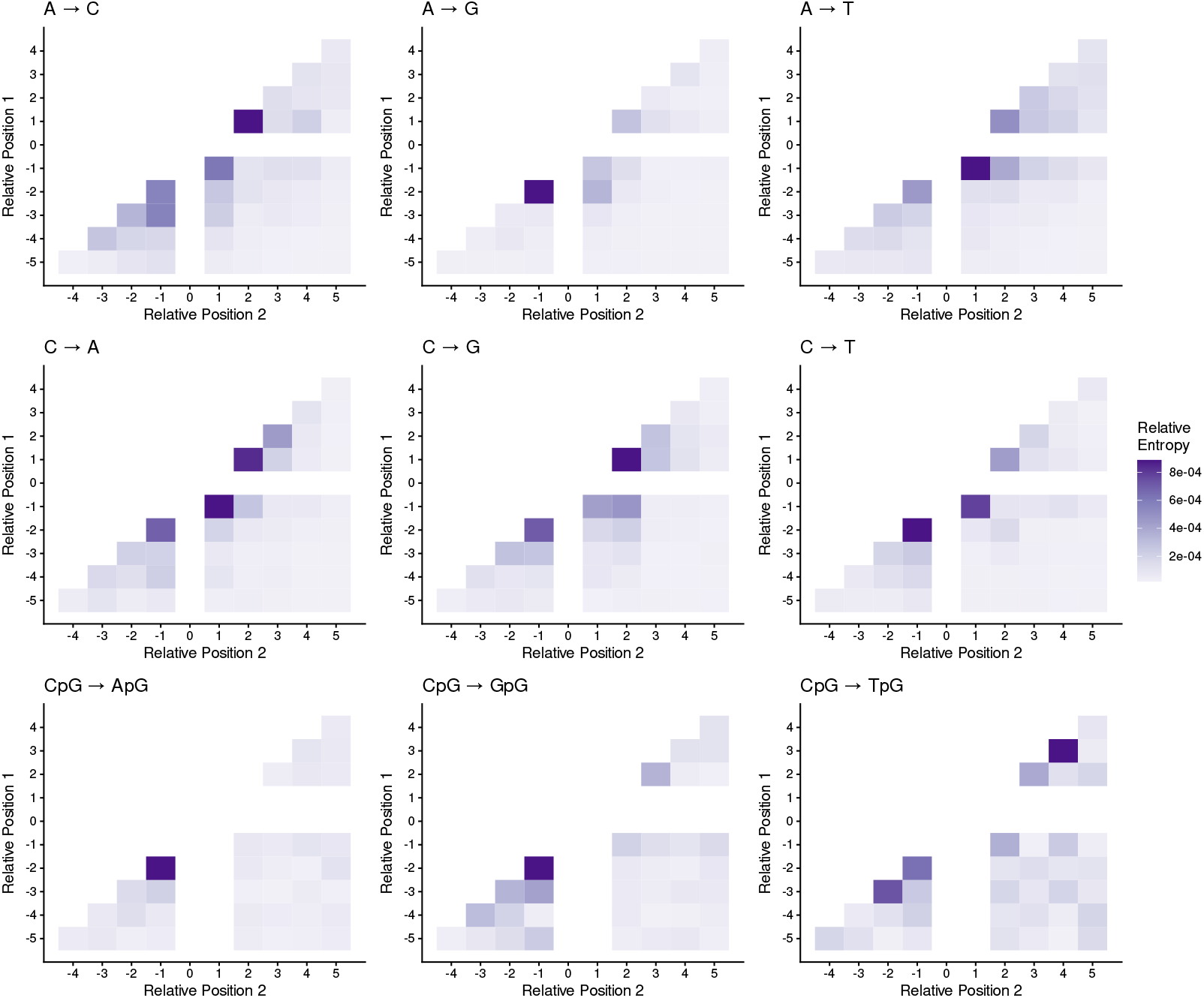
Two-position interaction effects on germline mutation rates.

**Fig. 7:**
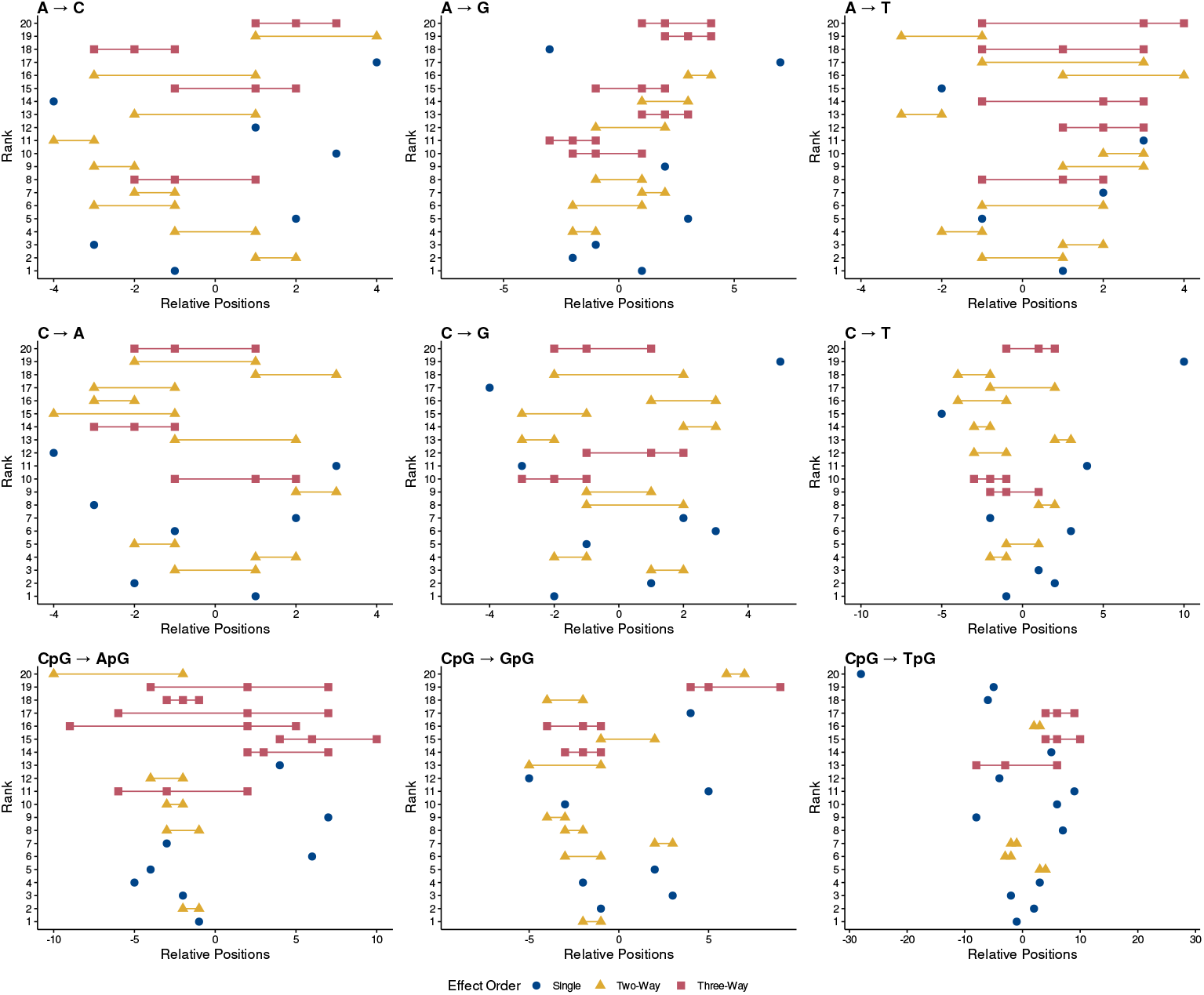
Rank hierarchy of top-ranked sequence context effects of single, two, and three-position effects. Top 25 single, two-position, and three-position effects ranked by RE. Across all subtypes, the top five highest-ranked terms are exclusively single- or two-position effects. For all non-CpG subtypes the strongest effects are concentrated within 10 bases of the focal position

**Fig. 8:**
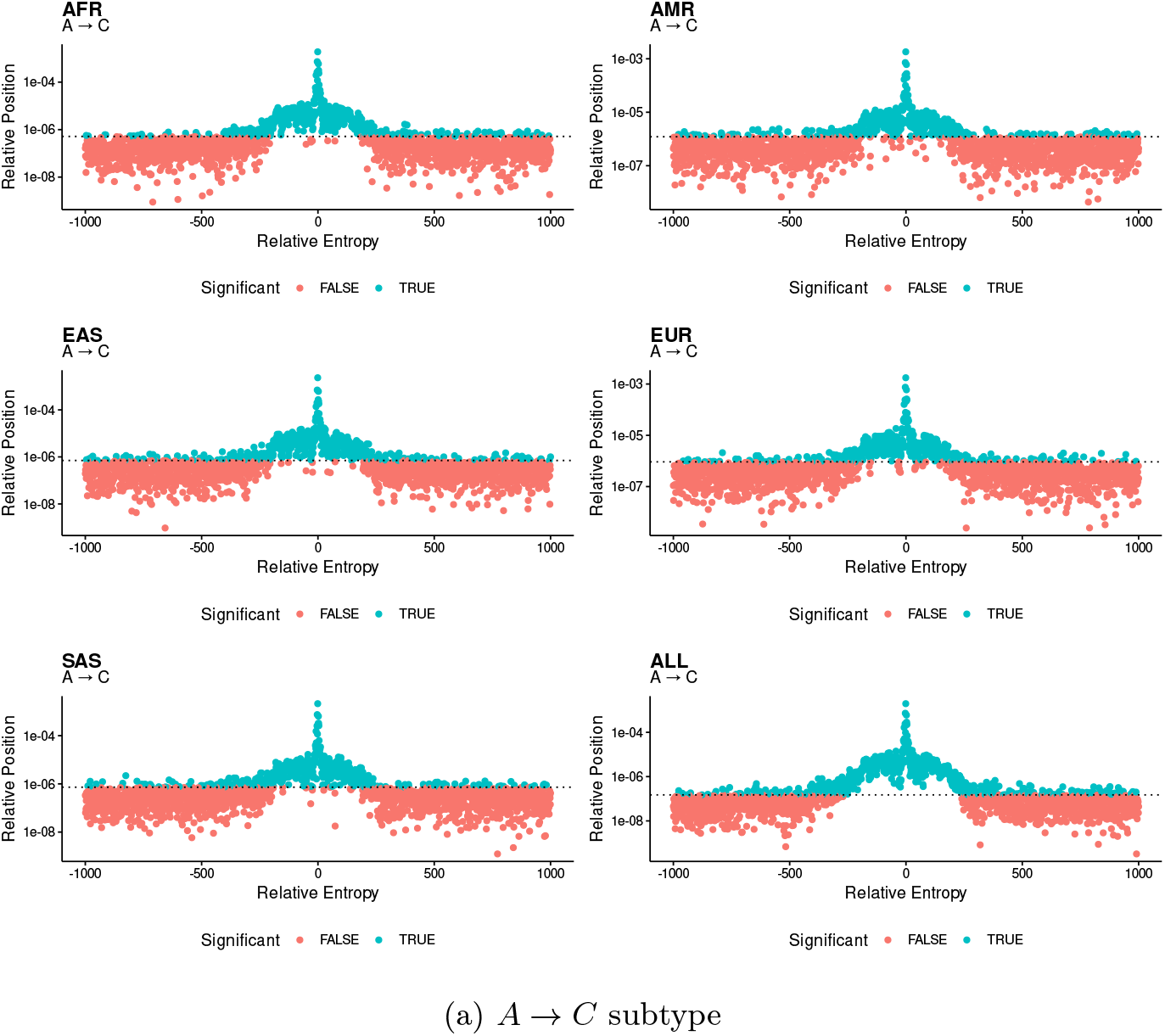
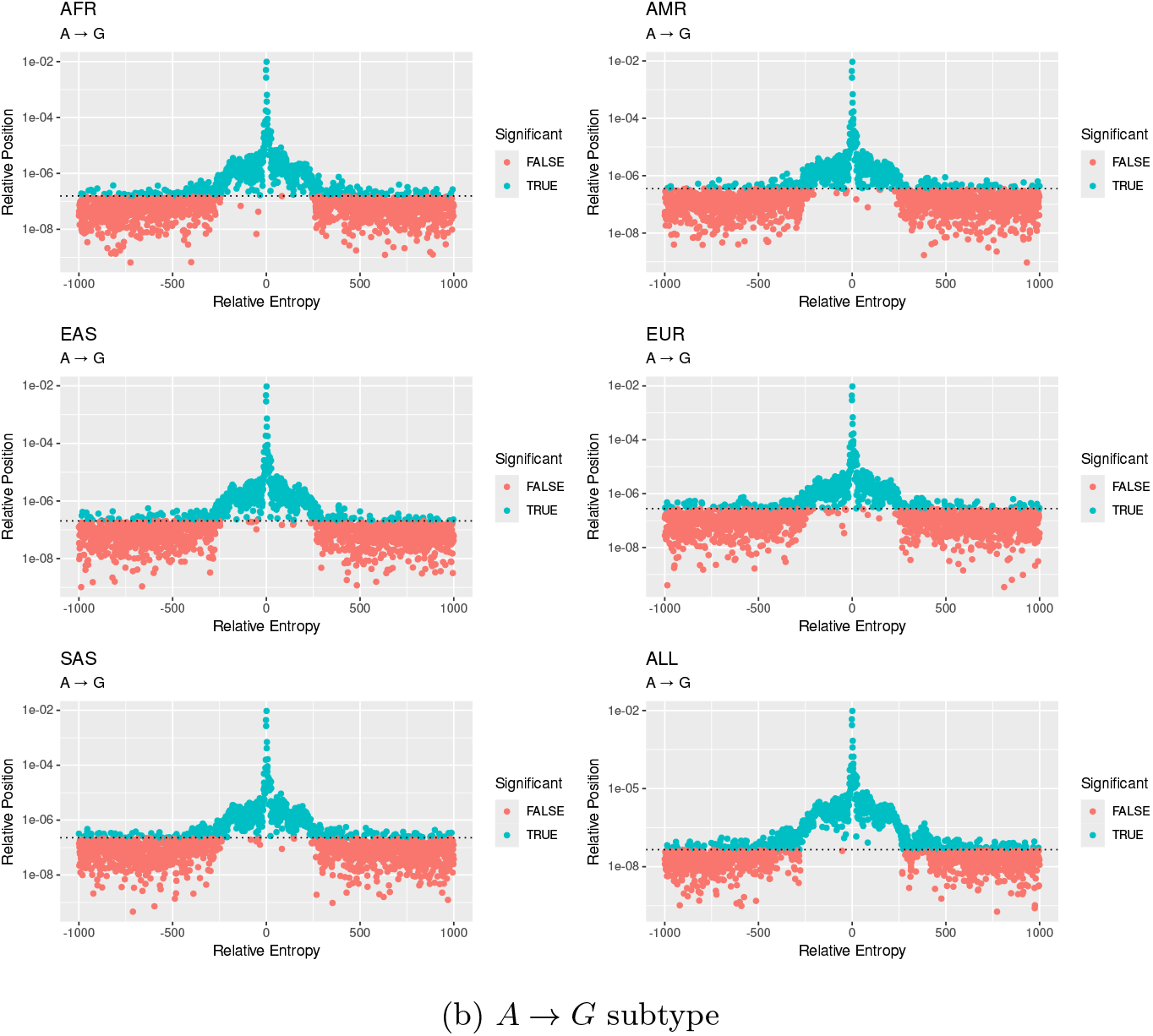
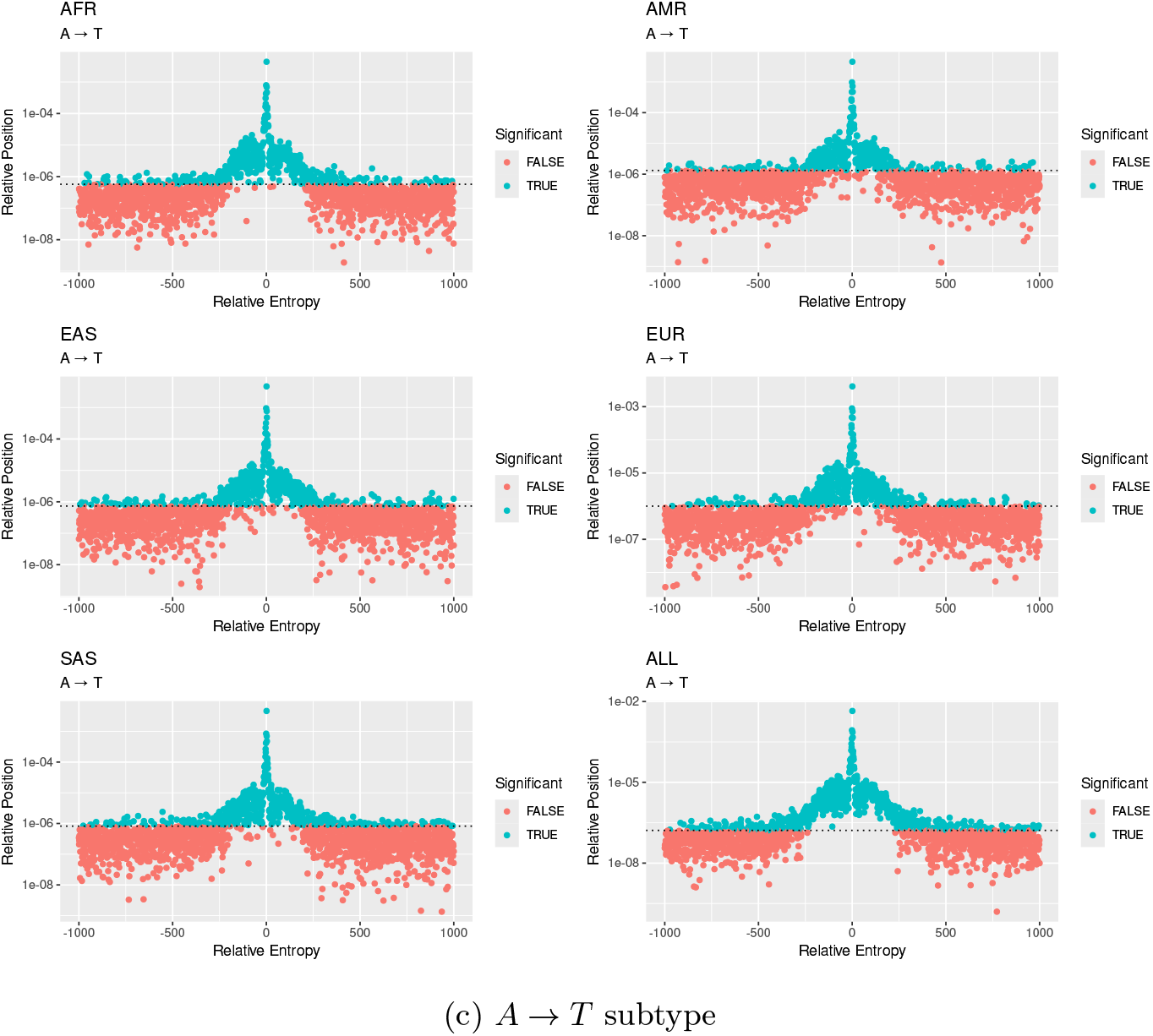
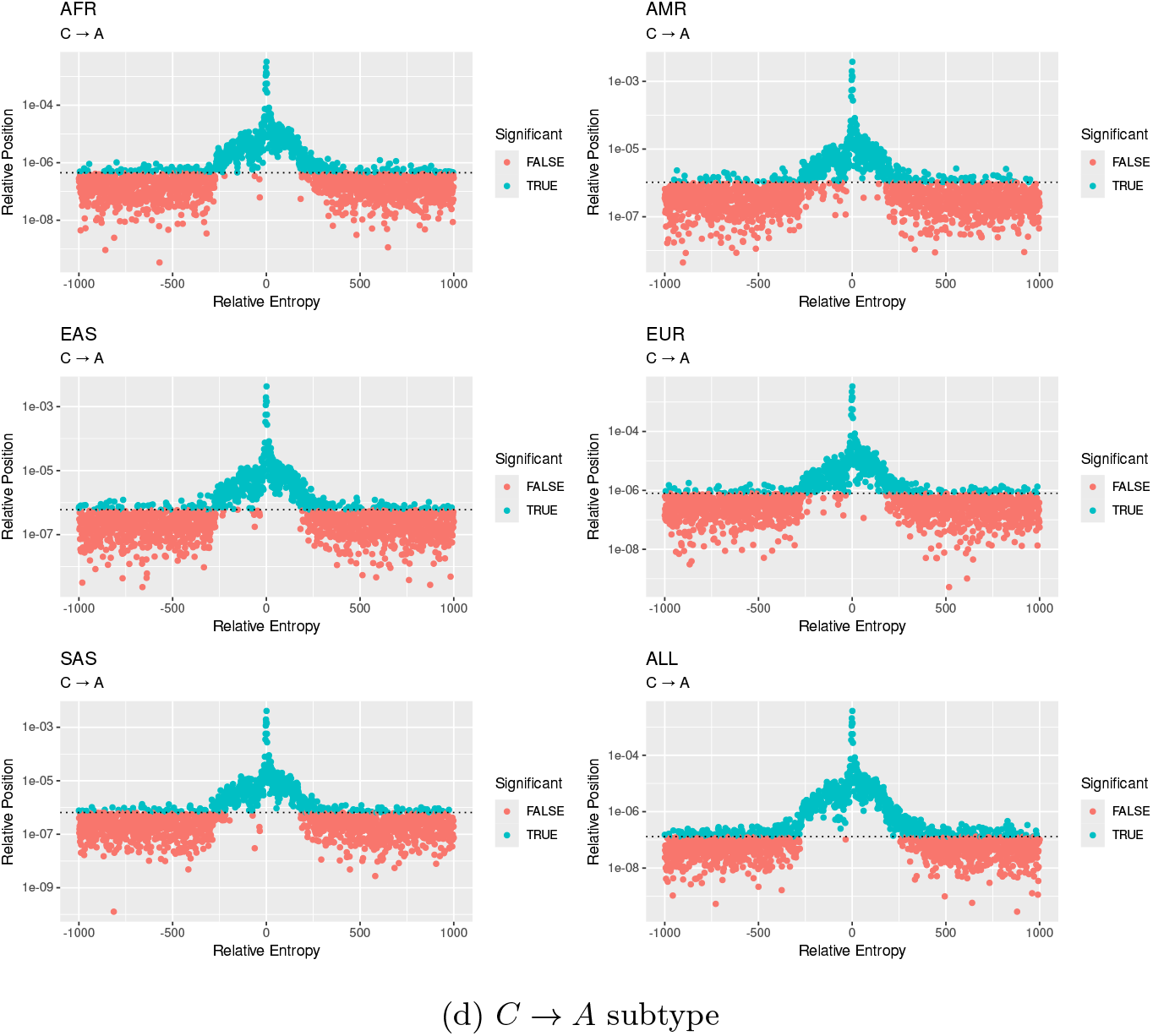
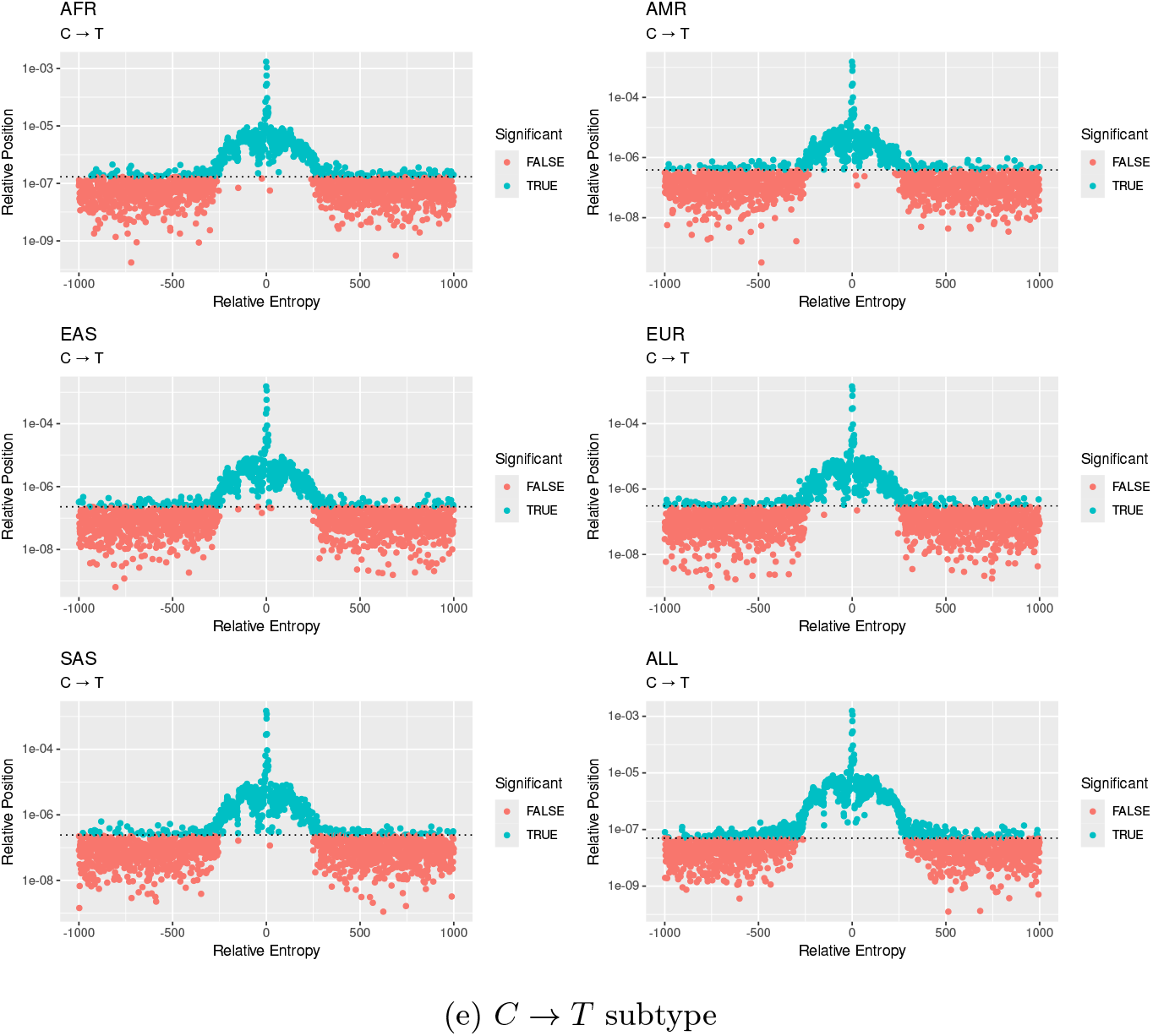
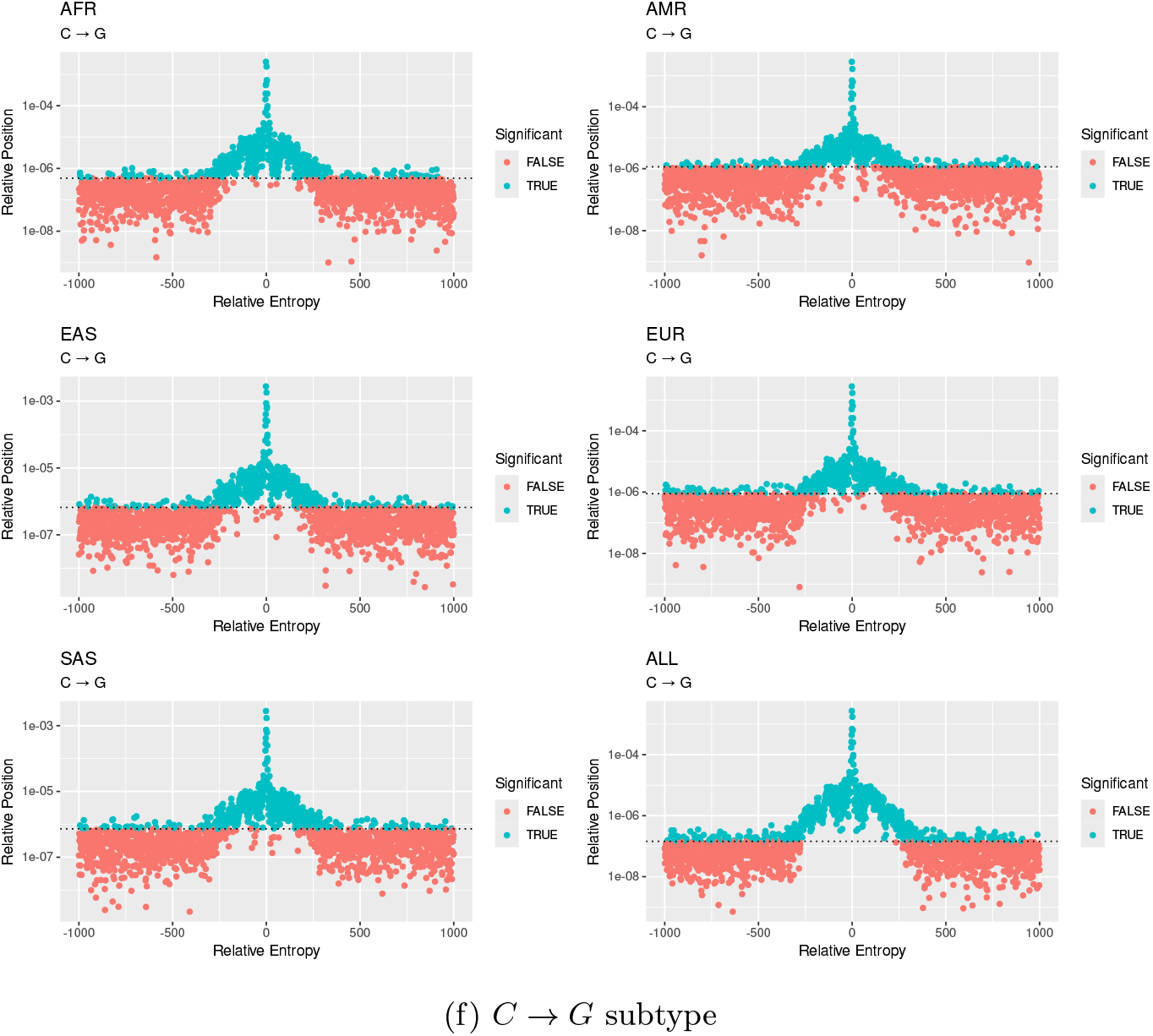
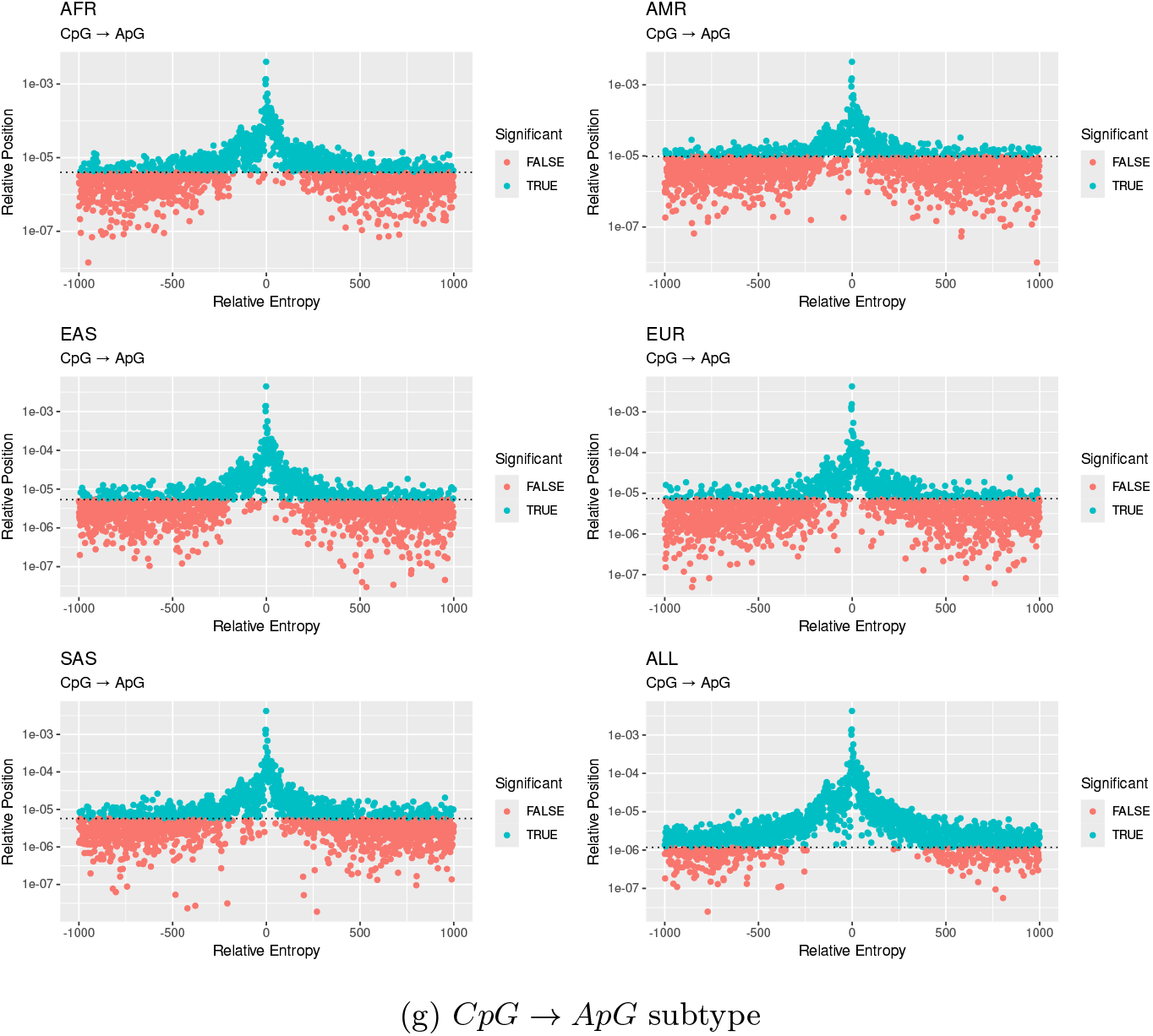
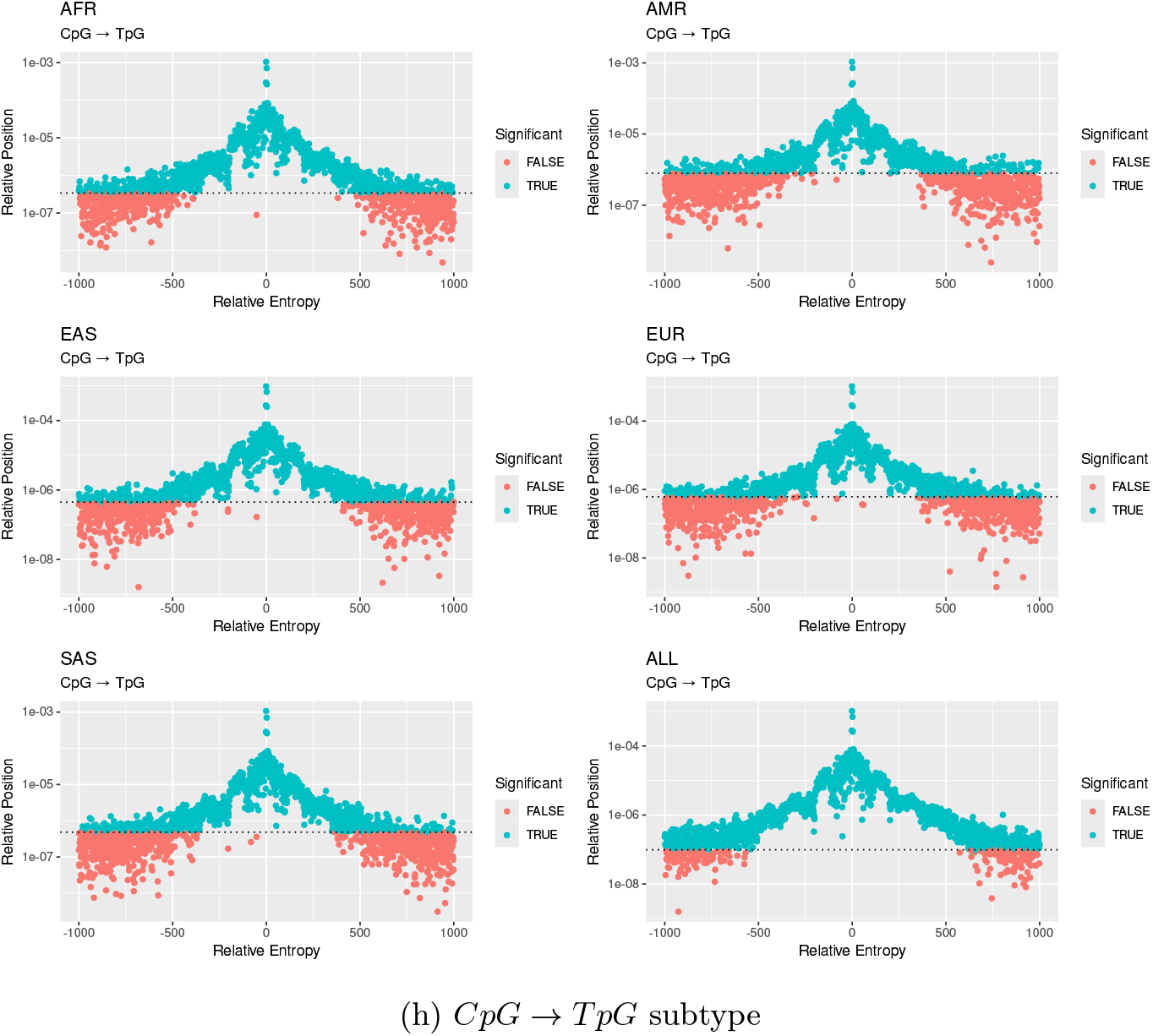
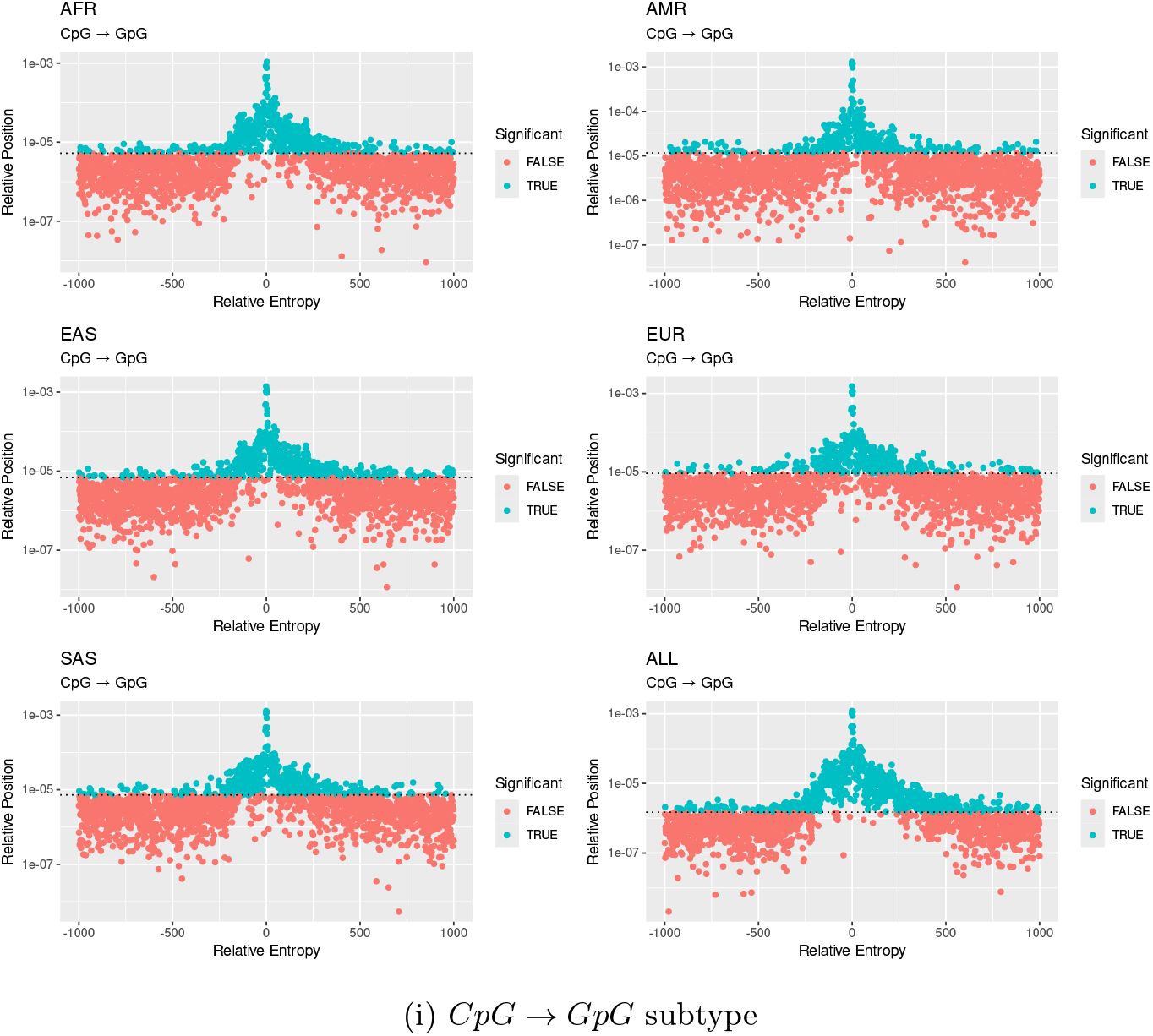
Population concordance of marginal influence of individual flanking positions across 1kGP populations.

**Fig. 9:**
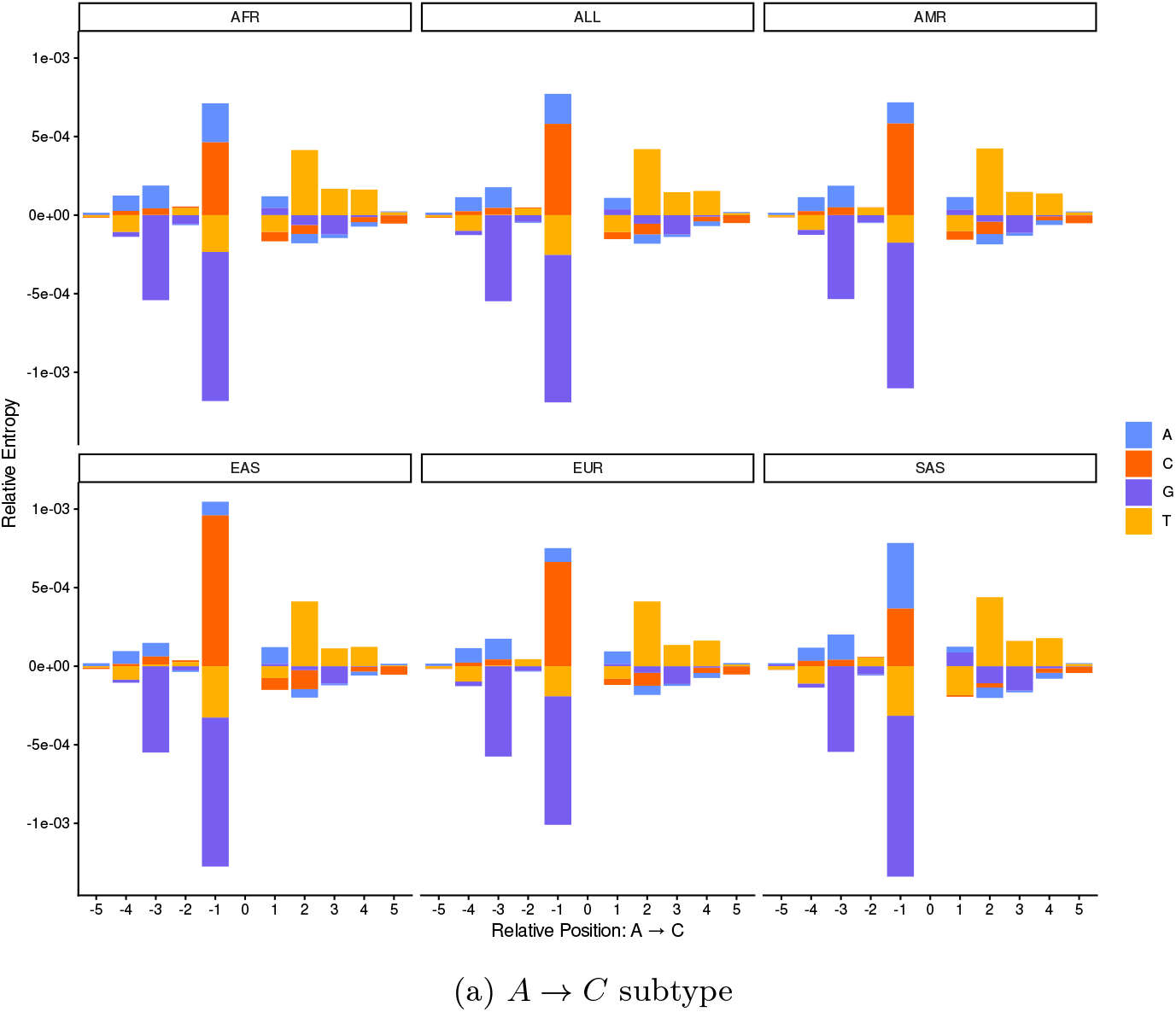
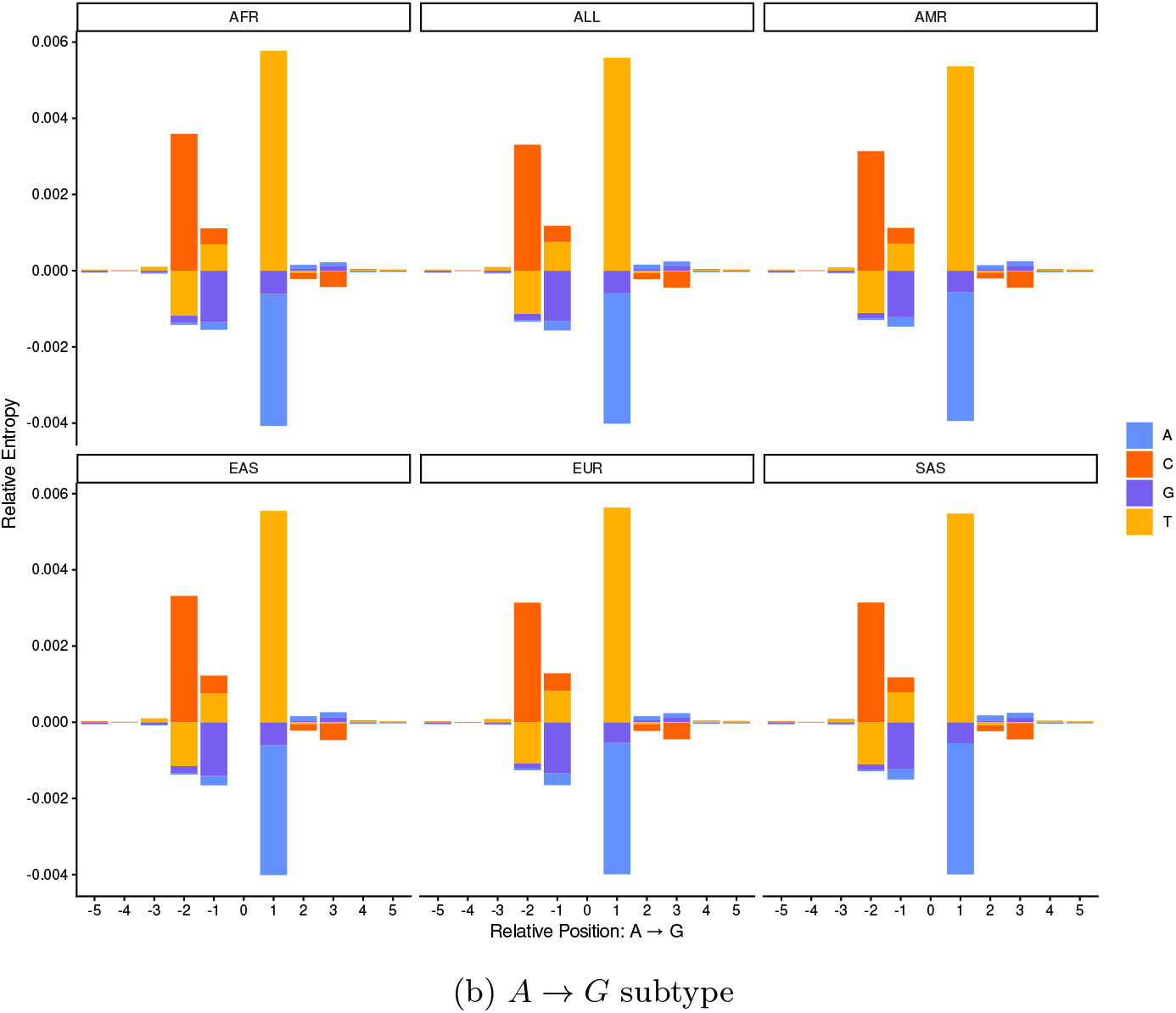
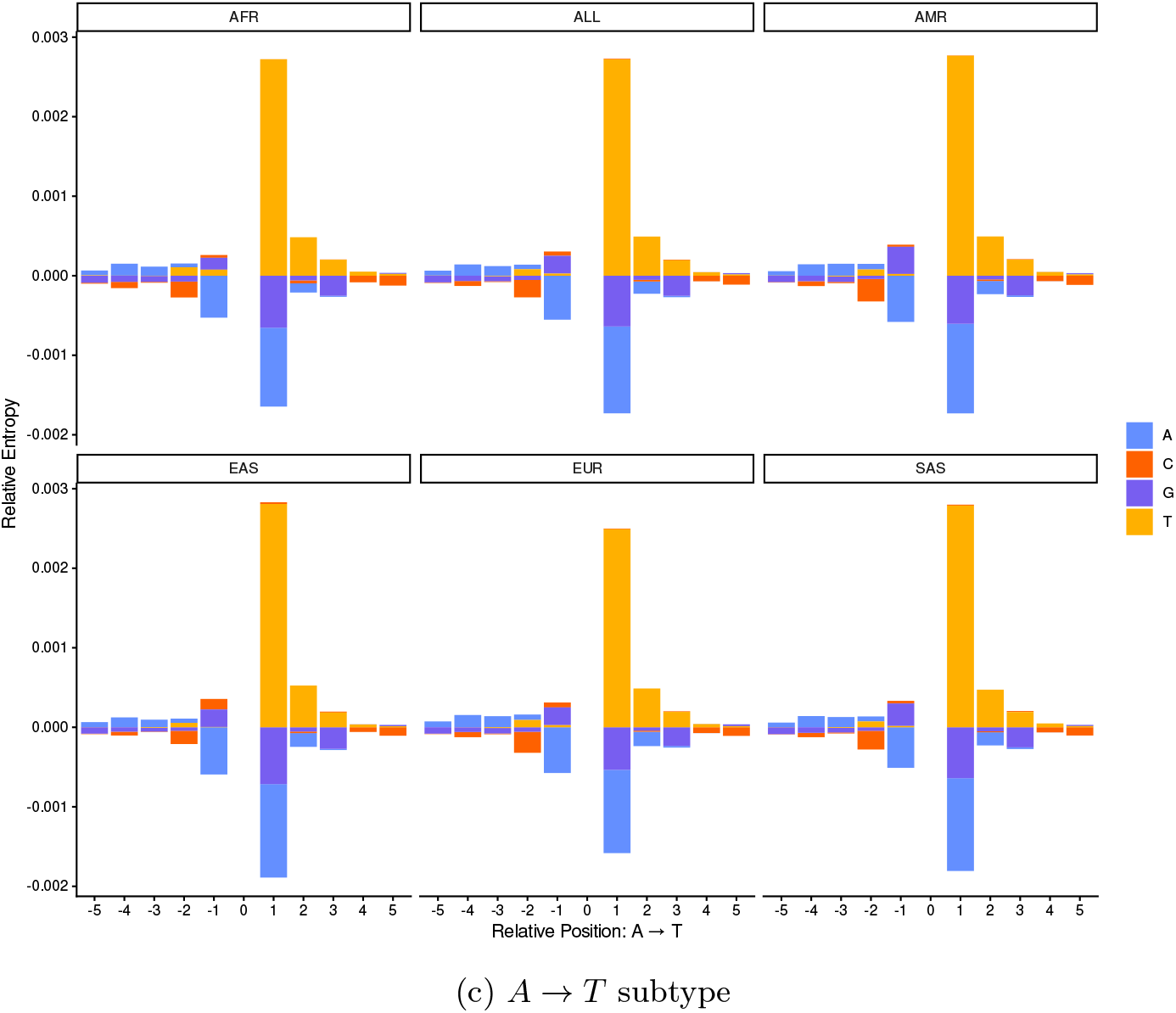
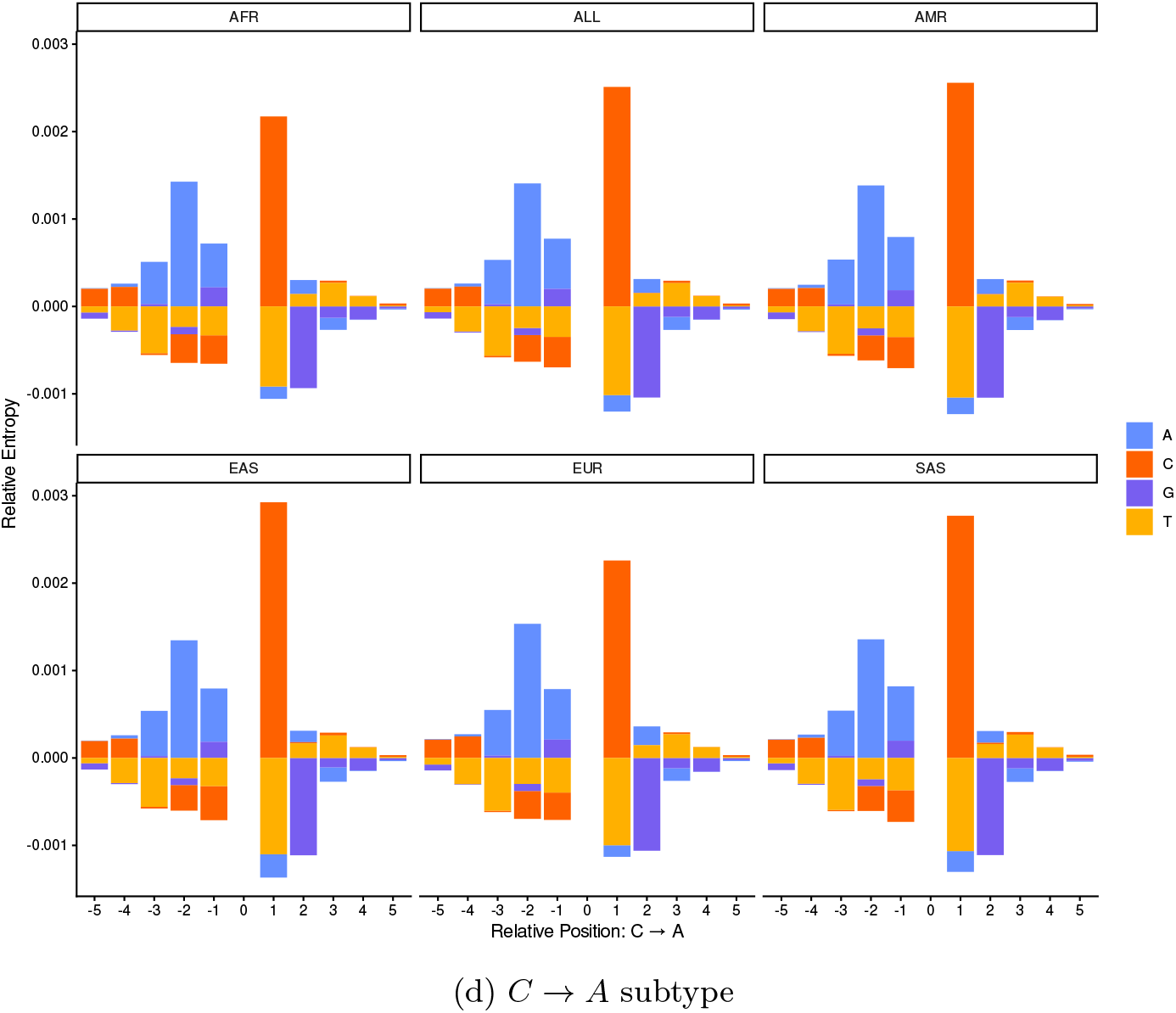
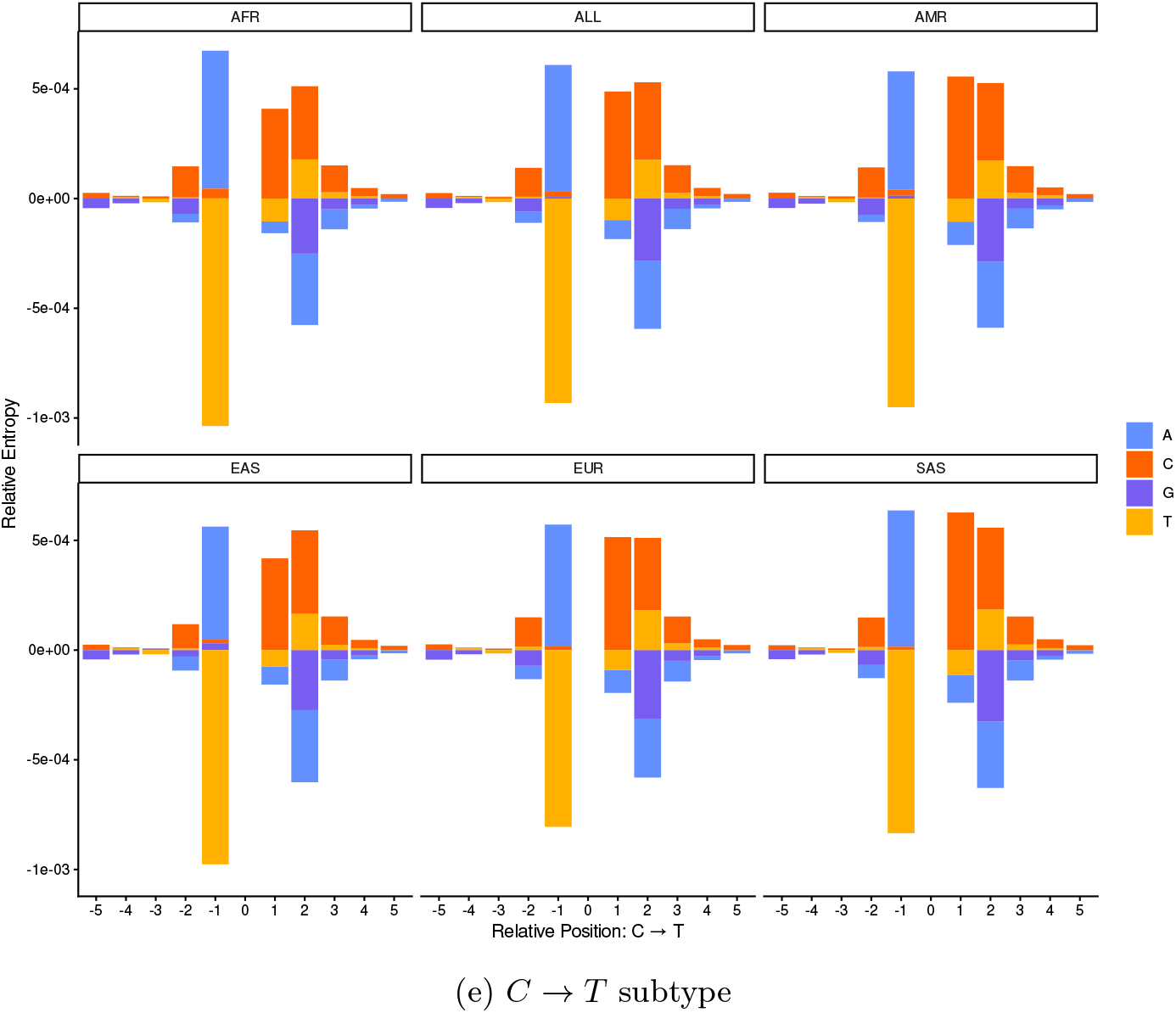
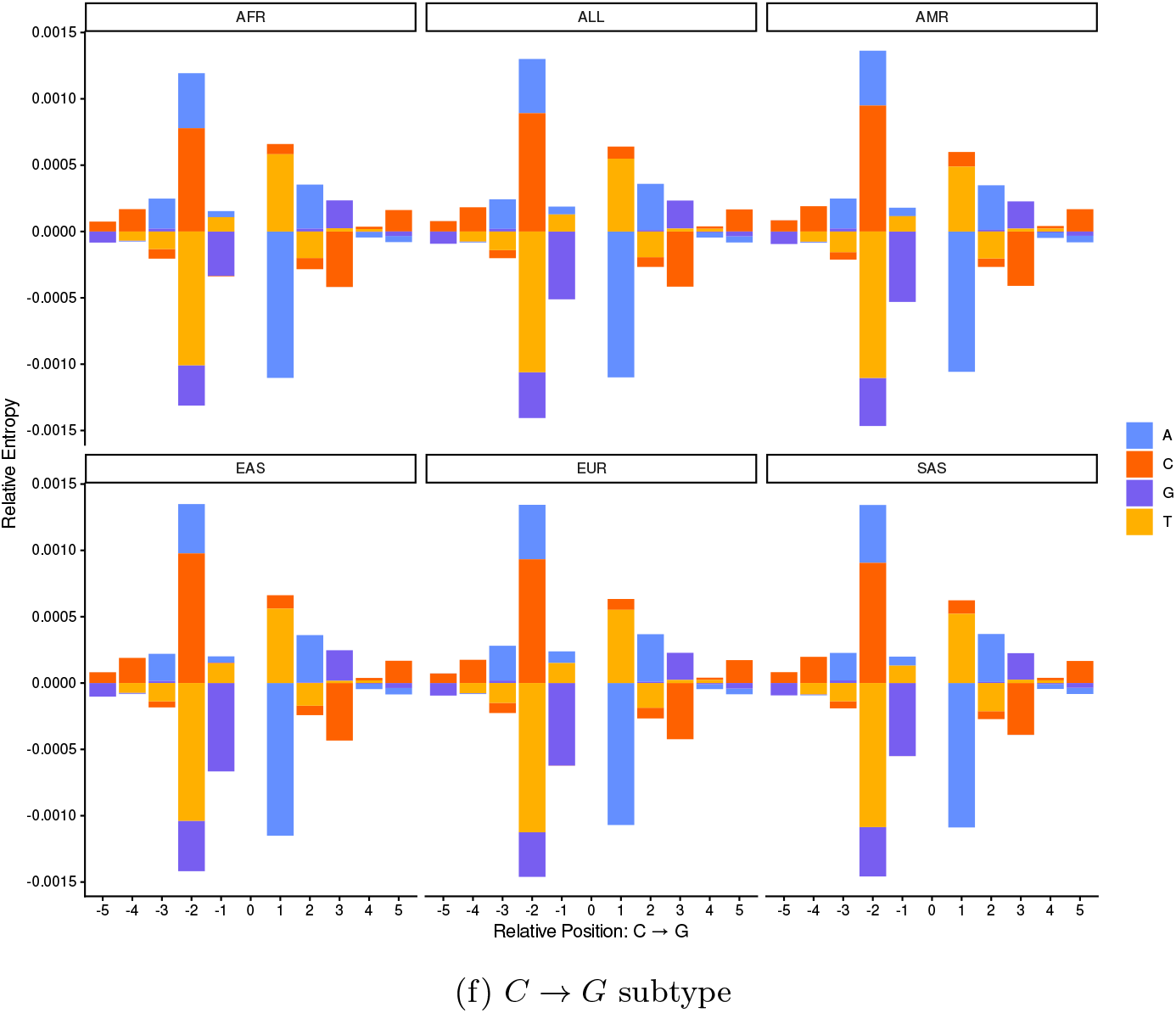
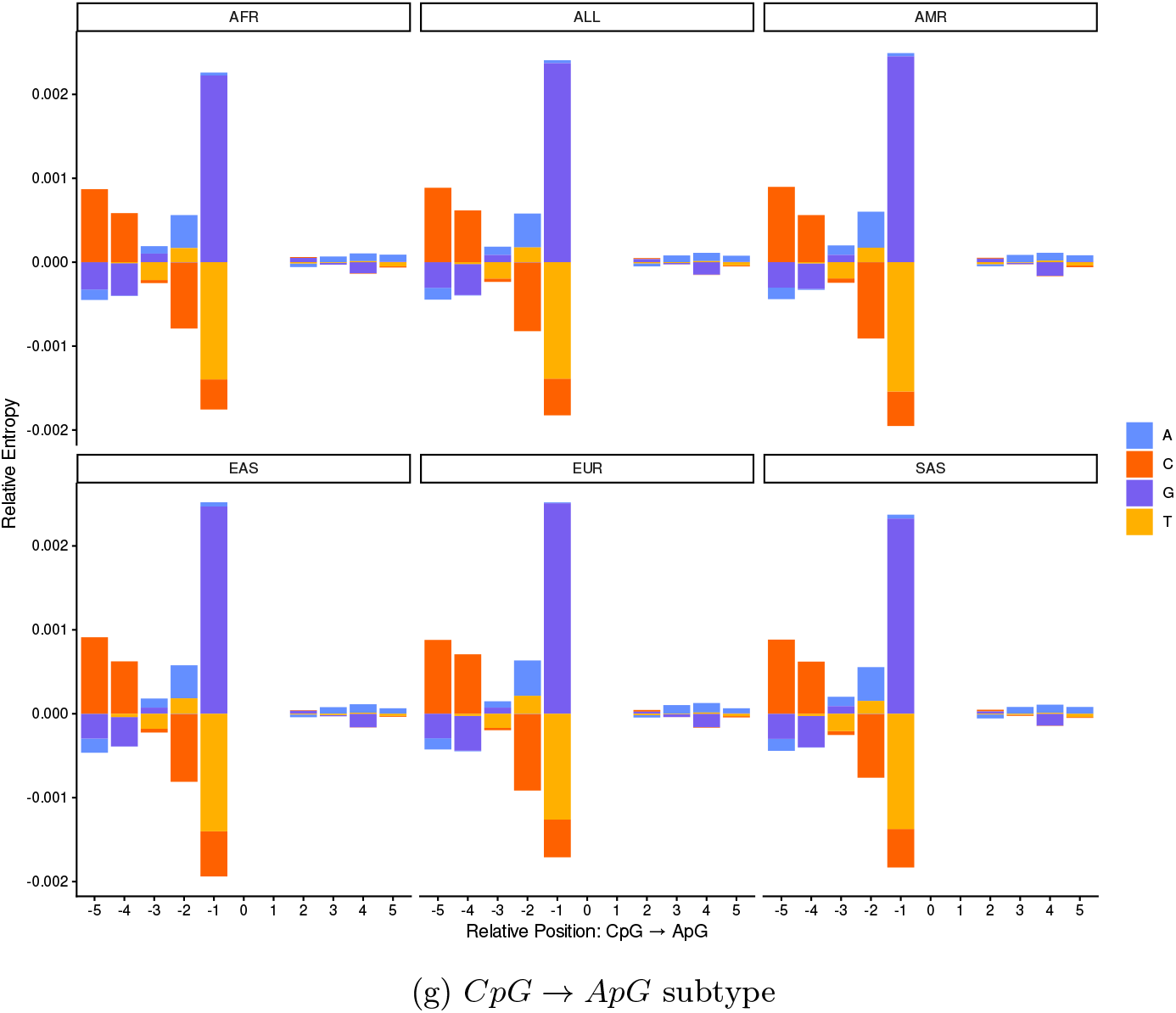
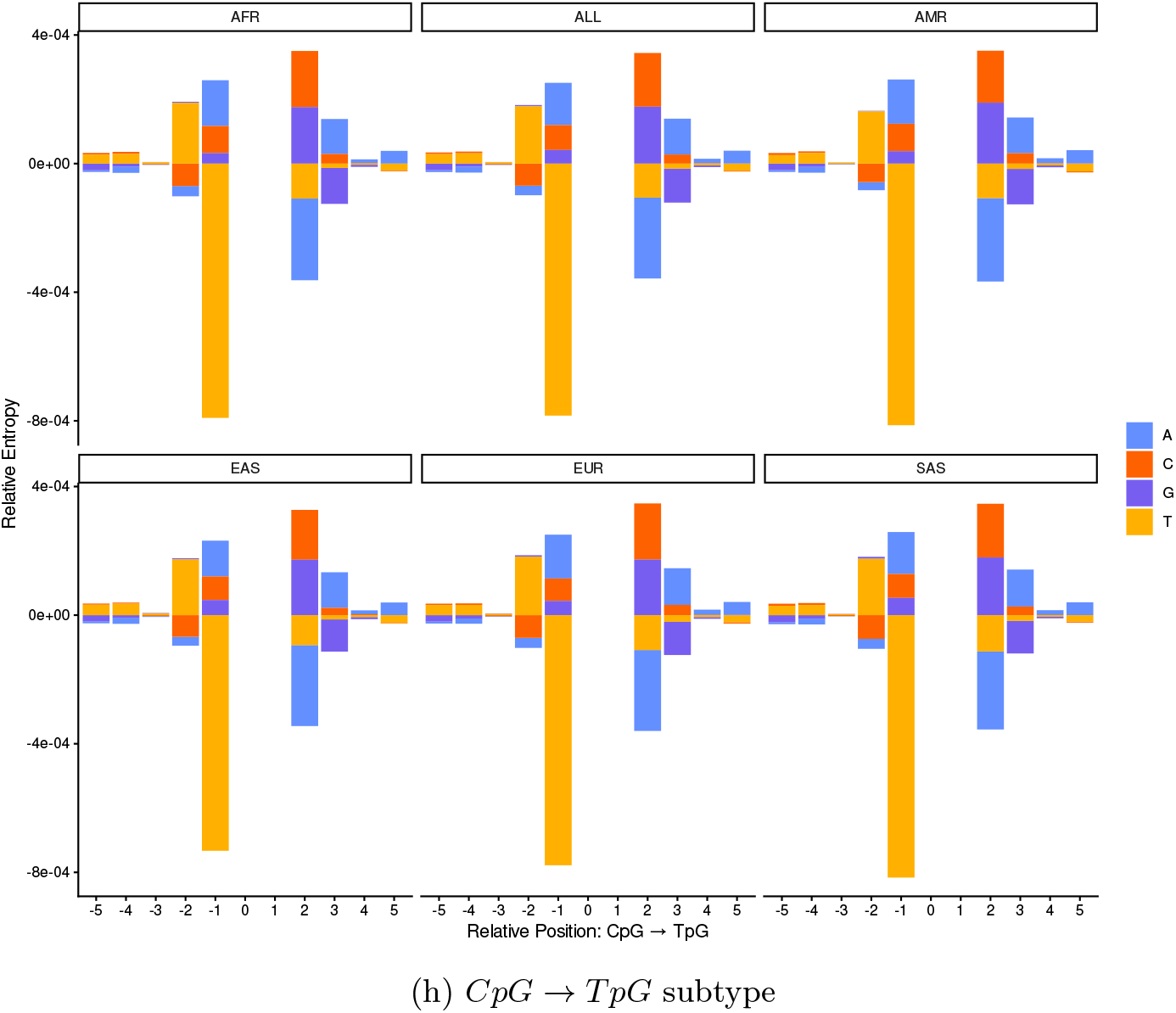
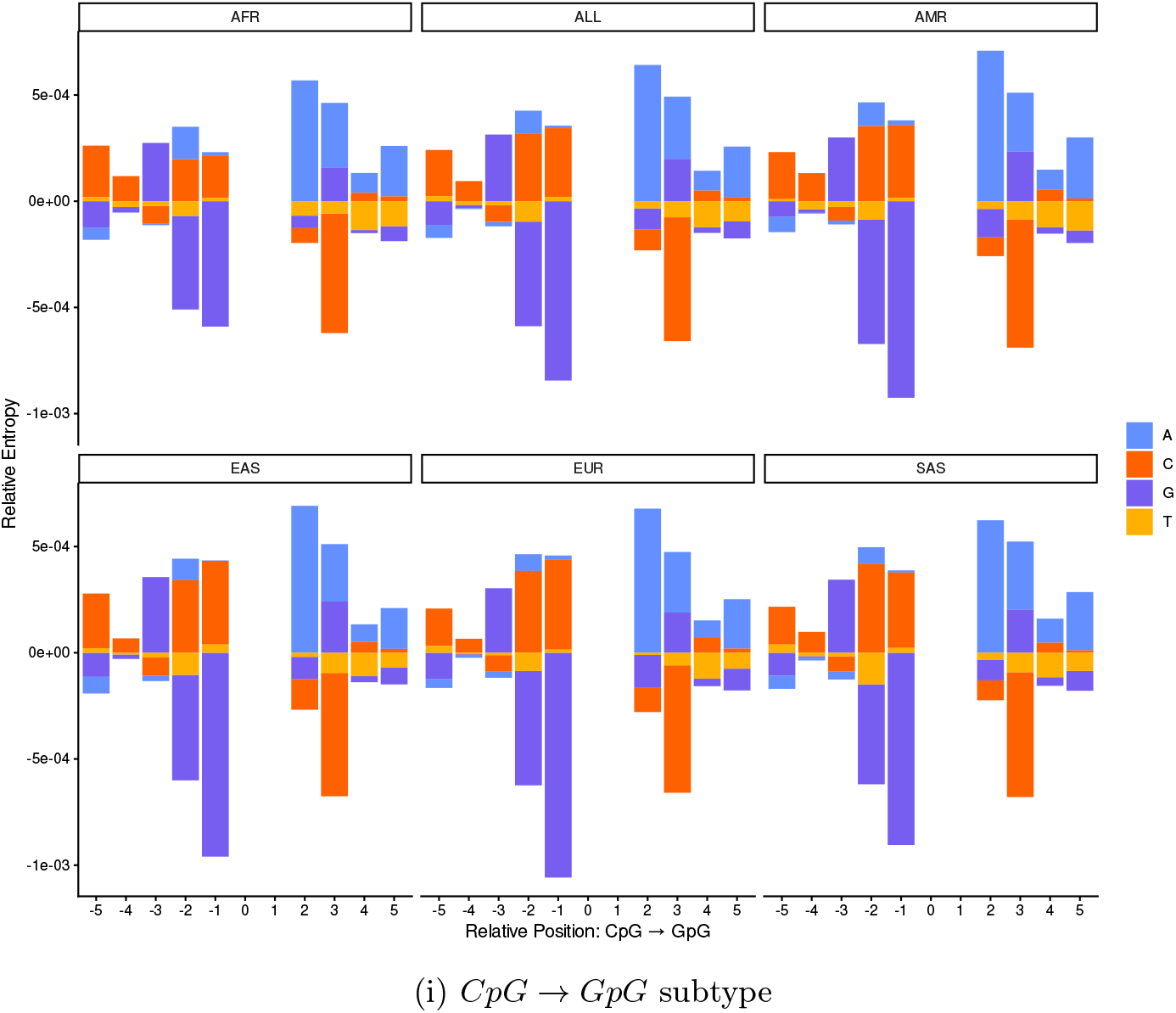
Population concordance of nucleotide-residuals in single position models across 1kGP groups.

## References

[1] Ségurel, L., Wyman, M.J., Przeworski, M.: Determinants of mutation rate variation in the human germline. Annual Review of Genomics and Human Genetics 15(1), 47–70 (2014) 10.1146/annurev-genom-031714-125740

[2] Loay, H., Kar, P., Koch, E., Seplyarskiy, V., Weghorn, D.: Mutation rate heterogeneity biases variant effect prediction and reveals genuine mutational robustness. The American Journal of Human Genetics 113(10), 2042–2054 (2026) 10.1016/j.ajhg.2026.08.011

[3] Rahbari, R., Wuster, A., Lindsay, S.J., Hardwick, R.J., Alexandrov, L.B., Al Turki, S., Dominiczak, A., Morris, A., Porteous, D., Smith, B., al.: Timing, rates and spectra of human germline mutation. Nature Genetics 48(2), 126–133 (2015) 10.1038/ng.3469

[4] Lercher, M.J., Hurst, L.D.: Human snp variability and mutation rate are higher in regions of high recombination. Trends in Genetics 18(7), 337–340 (2002) 10.1016/s0168-9525(02)02669-0

[5] Francioli, L.C., Polak, P.P., Koren, A., Menelaou, A., Chun, S., Renkens, I., Duijn, C.M., Swertz, M., Wijmenga, C., Ommen, G., al.: Genome-wide patterns and properties of de novo mutations in humans. Nature Genetics 47(7), 822–826 (2015) 10.1038/ng.3292

[6] Schuster-Böckler, B., Lehner, B.: Chromatin organization is a major influence on regional mutation rates in human cancer cells. Nature 488(7412), 504–507 (2012) 10.1038/nature11273

[7] Aggarwala, V., Voight, B.F.: An expanded sequence context model broadly explains variability in polymorphism levels across the human genome. Nature Genetics 48(4), 349–355 (2016) 10.1038/ng.3511

[8] Zhu, Y., Neeman, T., Yap, V.B., Huttley, G.A.: Statistical methods for identifying sequence motifs affecting point mutations. Genetics 205(2), 843–856 (2017) 10.1534/genetics.116.195677

[9] Carlson, J., Locke, A.E., Flickinger, M., Zawistowski, M., Levy, S., Myers, R.M., Boehnke, M., Kang, H.M., Scott, L.J., Li, J.Z., al.: Extremely rare variants reveal patterns of germline mutation rate heterogeneity in humans. Nature Communications 9(1) (2018) 10.1038/s41467-018-05936-5

[10] Seplyarskiy, V., Lee, D.J., Koch, E.M., Lichtman, J.S., Luan, H.H., Sunyaev, S.R.: A mutation rate model at the basepair resolution identifies the mutagenic effect of polymerase iii transcription (2022) 10.1101/2022.08.20.504670

[11] Byrska-Bishop, M., Evani, U.S., Zhao, X., Basile, A.O., Abel, H.J., Regier, A.A., Corvelo, A., Clarke, W.E., Musunuri, R., Nagulapalli, K., al.: High-coverage whole-genome sequencing of the expanded 1000 genomes project cohort including 602 trios. Cell 185(18) (2022) 10.1016/j.cell.2022.08.004

[12] Duncan, B.K., Miller, J.H.: Mutagenic deamination of cytosine residues in DNA. Nature 287(5782), 560–561 (1980)

[13] Alexandrov, L.B., Kim, J., Haradhvala, N.J., Huang, M.N., Tian Ng, A.W., Wu, Y., Boot, A., Covington, K.R., Gordenin, D.A., Bergstrom, E.N., Islam, S.M.A., Lopez-Bigas, N., Klimczak, L.J., McPherson, J.R., Morganella, S., Sabarinathan, R., Wheeler, D.A., Mustonen, V., Alexandrov, L.B., Bergstrom, E.N., Boot, A., Boutros, P., Chan, K., Covington, K.R., Fujimoto, A., Getz, G., Gordenin, D.A., Haradhvala, N.J., Huang, M.N., Islam, S.M.A., Kazanov, M., Kim, J., Klimczak, L.J., Lopez-Bigas, N., Lawrence, M., Martincorena, I., McPherson, J.R., Morganella, S., Mustonen, V., Nakagawa, H., Tian Ng, A.W., Polak, P., Prokopec, S., Roberts, S.A., Rozen, S.G., Sabarinathan, R., Saini, N., Shibata, T., Shiraishi, Y., Stratton, M.R., Teh, B.T., Vázquez-García, I., Wheeler, D.A., Wu, Y., Yousif, F., Yu, W., Getz, G., Rozen, S.G., Stratton, M.R., Aaltonen, L.A., Abascal, F., Abeshouse, A., Aburatani, H., Adams, D.J., Agrawal, N., Ahn, K.S., Ahn, S.-M., Aikata, H., Akbani, R., Akdemir, K.C., Al-Ahmadie, H., Al-Sedairy, S.T., Al-Shahrour, F., Alawi, M., Albert, M., Aldape, K., Alexandrov, L.B., Ally, A., Alsop, K., Alvarez, E.G., Amary, F., Amin, S.B., Aminou, B., Ammerpohl, O., Anderson, M.J., Ang, Y., Antonello, D., Anur, P., Aparicio, S., Appelbaum, E.L., Arai, Y., Aretz, A., Arihiro, K., Ariizumi, S.-i., Armenia, J., Arnould, L., Asa, S., Assenov, Y., Atwal, G., Aukema, S., Auman, J.T., Aure, M.R.R., Awadalla, P., Aymerich, M., Bader, G.D., Baez-Ortega, A., Bailey, M.H., Bailey, P.J., Balasundaram, M., Balu, S., Bandopadhayay, P., Banks, R.E., Barbi, S., Barbour, A.P., Barenboim, J., Barnholtz-Sloan, J., Barr, H., Barrera, E., Bartlett, J., Bartolome, J., Bassi, C., Bathe, O.F., Baumhoer, D., Bavi, P., Baylin, S.B., Bazant, W., Beardsmore, D., Beck, T.A., Behjati, S., Behren, A., Niu, B., Bell, C., Beltran, S., Benz, C., Berchuck, A., Bergmann, A.K., Bergstrom, E.N., Berman, B.P., Berney, D.M., Bernhart, S.H., Beroukhim, R., Berrios, M., Bersani, S., Bertl, J., Betancourt, M., Bhandari, V., Bhosle, S.G., Biankin, A.V., Bieg, M., Bigner, D., Binder, H., Birney, E., Birrer, M., Biswas, N.K., Bjerkehagen, B., Bodenheimer, T., Boice, L., Bonizzato, G., De Bono, J.S., Boot, A., Bootwalla, M.S., Borg, A., Borkhardt, A., Boroevich, K.A., Borozan, I., Borst, C., Bosenberg, M., Bosio, M., Boultwood, J., Bourque, G., Boutros, P.C., Bova, G.S., Bowen, D.T., Bowlby, R., Bowtell, D.D.L., Boyault, S., Boyce, R., Boyd, J., Brazma, A., Brennan, P., Brewer, D.S., Brinkman, A.B., Bristow, R.G., Broad-dus, R.R., Brock, J.E., Brock, M., Broeks, A., Brooks, A.N., Brooks, D., Brors, B., Brunak, S., Bruxner, T.J.C., Bruzos, A.L., Buchanan, A., Buchhalter, I., Buchholz, C., Bullman, S., Burke, H., Burkhardt, B., Burns, K.H., Busanovich, J., Bustamante, C.D., Butler, A.P., Butte, A.J., Byrne, N.J., Børresen-Dale, A.-L., Caesar-Johnson, S.J., Cafferkey, A., Cahill, D., Calabrese, C., Caldas, C., Calvo, F., Camacho, N., Campbell, P.J., Campo, E., Cantù, C., Cao, S., Carey, T.E., Carlevaro-Fita, J., Carlsen, R., Cataldo, I., Cazzola, M., Cebon, J., Cerfolio, R., Chadwick, D.E., Chakravarty, D., Chalmers, D., Chan, C.W.Y., Chan, K., Chan-Seng-Yue, M., Chandan, V.S., Chang, D.K., Chanock, S.J., Chantrill, L.A., Chateigner, A., Chatterjee, N., Chayama, K., Chen, H.-W., Chen, J., Chen, K., Chen, Y., Chen, Z., Cherniack, A.D., Chien, J., Chiew, Y.-E., Chin, S.-F., Cho, J., Cho, S., Choi, J.K., Choi, W., Chomienne, C., Chong, Z., Choo, S.P., Chou, A., Christ, A.N., Christie, E.L., Chuah, E., Cibulskis, C., Cibulskis, K., Cingarlini, S., Clapham, P., Claviez, A., Cleary, S., Cloonan, N., Cmero, M., Collins, C.C., Connor, A.A., Cooke, S.L., Cooper, C.S., Cope, L., Corbo, V., Cordes, M.G., Cordner, S.M., Cortés-Ciriano, I., Covington, K., Cowin, P.A., Craft, B., Craft, D., Creighton, C.J., Cun, Y., Curley, E., Cutcutache, I., Czajka, K., Czerniak, B., Dagg, R.A., Danilova, L., Davi, M.V., Davidson, N.R., Davies, H., Davis, I.J., Davis-Dusenbery, B.N., Dawson, K.J., De La Vega, F.M., De Paoli-Iseppi, R., Defreitas, T., Tos, A.P.D., Delaneau, O., Demchok, J.A., Demeulemeester, J., Demidov, G.M., Demircioğlu, D., Dennis, N.M., Denroche, R.E., Dentro, S.C., Desai, N., Deshpande, V., Deshwar, A.G., Desmedt, C., Deu-Pons, J., Dhalla, N., Dhani, N.C., Dhingra, P., Dhir, R., DiBiase, A., Diamanti, K., Ding, L., Ding, S., Dinh, H.Q., Dirix, L., Doddapaneni, H., Donmez, N., Dow, M.T., Drapkin, R., Drechsel, O., Drews, R.M., Serge, S., Dudderidge, T., Dueso-Barroso, A., Dunford, A.J., Dunn, M., Dursi, L.J., Duthie, F.R., Dutton-Regester, K., Eagles, J., Easton, D.F., Edmonds, S., Edwards, P.A., Edwards, S.E., Eeles, R.A., Ehinger, A., Eils, J., Eils, R., El-Naggar, A., Eldridge, M., Ellrott, K., Erkek, S., Escaramis, G., Espiritu, S.M.G., Estivill, X., Etemadmoghadam, D., Eyfjord, J.E., Faltas, B.M., Fan, D., Fan, Y., Faquin, W.C., Farcas, C., Fassan, M., Fatima, A., Favero, F., Fayzullaev, N., Felau, I., Fereday, S., Ferguson, M.L., Ferretti, V., Feuerbach, L., Field, M.A., Fink, J.L., Finocchiaro, G., Fisher, C., Fittall, M.W., Fitzgerald, A., Fitzgerald, R.C., Flanagan, A.M., Fleshner, N.E., Flicek, P., Foekens, J.A., Fong, K.M., Fonseca, N.A., Foster, C.S., Fox, N.S., Fraser, M., Frazer, S., Frenkel-Morgenstern, M., Friedman, W., Frigola, J., Fronick, C.C., Fujimoto, A., Fujita, M., Fukayama, M., Fulton, L.A., Fulton, R.S., Furuta, M., Futreal, P.A., Füllgrabe, A., Gabriel, S.B., Gallinger, S., Gambacorti-Passerini, C., Gao, J., Gao, S., Garraway, L Garred, Garrison, E., Garsed, D.W., Gehlenborg, N., Gelpi, J.L.L., George, J., Gerhard, D.S., Gerhauser, C., Gershenwald, J.E., Gerstein, M., Gerstung, M., Getz, G., Ghori, M., Ghossein, R., Giama, N.H., Gibbs, R.A., Gibson, B., Gill, A.J., Gill, P., Giri, D.D., Glodzik, D., Gnanapragasam, V.J., Goebler, M.E., Goldman, M.J., Gomez, C., Gonzalez, S., Gonzalez-Perez, A., Gordenin, D.A., Gossage, J., Gotoh, K., Govindan, R., Grabau, D., Graham, J.S., Grant, R.C., Green, A.R., Green, E., Greger, L., Grehan, N., Grimaldi, S., Grimmond, S.M., Grossman, R.L., Grundhoff, A., Gundem, G., Guo, Q., Gupta, M., Gupta, S., Gut, I.G., Gut, M., Göke, J., Ha, G., Haake, A., Haan, D., Haas, S., Haase, K., Haber, J.E., Habermann, N., Hach, F., Haider, S., Hama, N., Hamdy, F.C., Hamilton, A., Hamilton, M.P., Han, L., Hanna, G.B., Hansmann, M., Haradhvala, N.J., Harismendy, O., Harliwong, I., Harmanci, A.O., Harrington, E., Hasegawa, T., Haussler, D., Hawkins, S., Hayami, S., Hayashi, S., Hayes, D.N., Hayes, S.J., Hayward, N.K., Hazell, S., He, Y., Heath, A.P., Heath, S.C., Hedley, D., Hegde, A.M., Heiman, D.I., Heinold, M.C., Heins, Z., Heisler, L.E., Hellstrom-Lindberg, E., Helmy, M., Heo, S.G., Hepperla, A.J., Heredia-Genestar, J.M., Herrmann, C., Hersey, P., Hess, J.M., Hilmarsdottir, H., Hinton, J., Hirano, S., Hiraoka, N., Hoadley, K.A., Hobolth, A., Hodzic, E., Hoell, J.I., Hoffmann, S., Hofmann, O., Holbrook, A., Holik, A.Z., Hollingsworth, M.A., Holmes, O., Holt, R.A., Hong, C., Hong, E.P., Hong, J.H., Hooijer, G.K., Hornshøj, H., Hosoda, F., Hou, Y., Hovestadt, V., Howat, W., Hoyle, A.P., Hruban, R.H., Hu, J., Hu, T., Hua, X., Huang, K.-l., Huang, M., Huang, M.N., Huang, V., Huang, Y., Huber, W., Hudson, T.J., Hummel, M., Hung, J.A., Huntsman, D., Hupp, T.R., Huse, J., Huska, M.R., Hutter, B., Hutter, C.M., Hübschmann, D., Iacobuzio-Donahue, C.A., Imbusch, C.D., Imielinski, M., Imoto, S., Isaacs, W.B., Isaev, K., Ishikawa, S., Iskar, M., Islam, S.M.A., Ittmann, M., Ivkovic, S., Izarzugaza, J.M.G., Jacquemier, J., Jakrot, V., Jamieson, N.B., Jang, G.H., Jang, S.J., Jayaseelan, J.C., Jayasinghe, R., Jefferys, S.R., Jegalian, K., Jennings, J.L., Jeon, S.-H., Jerman, L., Ji, Y., Jiao, W., Johansson, P.A., Johns, A.L., Johns, J., Johnson, R., Johnson, T.A., Jolly, C., Joly, Y., Jonasson, J.G., Jones, C.D., Jones, D.R., Jones, D.T.W., Jones, N., Jones, S.J.M., Jonkers, J., Ju, Y.S., Juhl, H., Jung, J., Juul, M., Juul, R.I., Juul, S., Jäger, N., Kabbe, R., Kahles, A., Kahraman, A., Kaiser, V.B., Kakavand, H., Kalimuthu, S., Kalle, C., Kang, K.J., Karaszi, K., Karlan, B., Karlić, R., Karsch, D., Kasaian, K., Kassahn, K.S., Katai, H., Kato, M., Katoh, H., Kawakami, Y., Kay, J.D., Kazakoff, S.H., Kazanov, M.D., Keays, M., Kebebew, E., Kefford, R.F., Kellis, M., Kench, J.G., Kennedy, C.J., Kerssemakers, J.N.A., Khoo, D., Khoo, V., Khuntikeo, N., Khurana, E., Kilpinen, H., Kim, H.K., Kim, H.-L., Kim, H.-Y., Kim, H., Kim, J., Kim, J., Kim, J.K., Kim, Y., King, T.A., Klapper, W., Kleinheinz, K., Klimczak, L.J., Knappskog, S., Kneba, M., Knoppers, B.M., Koh, Y., Komorowski, J., Komura, D., Komura, M., Kong, G., Kool, M., Korbel, J.O., Korchina, V., Korshunov, A., Koscher, M., Koster, R., Kote-Jarai, Z., Koures, A., Kovacevic, M., Kremeyer, B., Kretzmer, H., Kreuz, M., Krishnamurthy, S., Kube, D., Kumar, K., Kumar, P., Kumar, S., Kumar, Y., Kundra, R., Kübler, K., Küppers, R., Lagergren, J., Lai, P.H., Laird, P.W., Lakhani, S.R., Lalansingh, C.M., Lalonde, E., Lamaze, F.C., Lambert, A., Lander, E., Landgraf, P., Landoni, L., Langerød, A., Lanzós, A., Larsimont, D., Larsson, E., Lathrop, M., Lau, L.M.S., Lawerenz, C., Lawlor, R.T., Lawrence, M.S., Lazar, A.J., Lazic, A.M., Le, X., Lee, D., Lee, D., Lee, E.A., Lee, H.J., Lee, J.J.-K., Lee, J.-Y., Lee, J., Lee, M.T.M., Lee-Six, H., Lehmann, K.-V., Lehrach, H., Lenze, D., Leonard, C.R., Leongamornlert, D.A., Leshchiner, I., Letourneau, L., Letunic, I., Levine, D.A., Lewis, L., Ley, T., Li, C., Li, C.H., Li, H.I., Li, J., Li, L., Li, S., Li, S., Li, X., Li, X., Li, X., Li, Y., Liang, H., Liang, S.-B., Lichter, P., Lin, P., Lin, Z., Linehan, W.M., Lingjærde, O.C., Liu, D., Liu, E.M., Liu, F.-F.F., Liu, F., Liu, J., Liu, X., Livingstone, J., Livitz, D., Livni, N., Lochovsky, L., Loeffler, M., Long, G.V., Lopez-Guillermo, A., Lou, S., Louis, D.N., Lovat, L.B., Lu, Y., Lu, Y.-J., Lu, Y., Luchini, C., Lungu, I., Luo, X., Luxton, H.J., Lynch, A.G., Lype, L., López, C., López-Otín, C., Ma, E.Z., Ma, Y., MacGrogan, G., MacRae, S., Macintyre, G., Madsen, T., Maejima, K., Mafficini, A., Maglinte, D.T., Maitra, A., Majumder, P.P., Malcovati, L., Mali-kic, S., Malleo, G., Mann, G.J., Mantovani-Löffler, L., Marchal, K., Marchegiani, G., Mardis, E.R., Margolin, A.A., Marin, M.G., Markowetz, F., Markowski, J., Marks, J., Marques-Bonet, T., Marra, M.A., Marsden, L., Martens, J.W.M., Martin, S., Martin-Subero, J.I., Martincorena, I., Martinez-Fundichely, A., Maruvka, Y.E., Mashl, R.J., Massie, C.E., Matthew, T.J., Matthews, L., Mayer, E., Mayes, S., Mayo, M., Mbabaali, F., McCune, K., McDermott, U., McGillivray, P.D., McLellan, M.D., McPherson, J.D., McPherson, J.R., McPherson, T.A., Meier, S.R., Meng, A., Meng, S., Menzies, A., Merrett, N.D., Merson, S., Meyerson, M., Meyerson, W., Mieczkowski, P.A., Mihaiescu, G.L., Mijalkovic, S., Mikkelsen, T., Milella, M., Mileshkin, L., Miller, C.A., Miller, D.K., Miller, J.K., Mills, G.B., Milovanovic, A., Minner, S., Miotto, M., Arnau, G.M., Mirabello, L., Mitchell, C., Mitchell, T.J., Miyano, S., Miyoshi, N., Mizuno, S., Molnár-Gábor, F., Moore, M.J., Moore, R.A., Morganella, S., Morris, Q.D., Morrison, C., Mose, L.E., Moser, C.D., Muiños, F., Mularoni, L., Mungall, A.J., Mungall, K., Musgrove, E.A., Mustonen, V., Mutch, D., Muyas, F., Muzny, D.M., Muñoz, A., Myers, J., Myklebost, O., Möller, P., Nagae, G., Nagrial, A.M., Nahal-Bose, H.K., Nakagama, H., Nakagawa, H., Nakamura, H., Nakamura, T., Nakano, K., Nandi, T., Nangalia, J., Nastic, M., Navarro, A., Navarro, F.C.P., Neal, D.E., Nettekoven, G., Newell, F., Newhouse, S.J., Newton, Y., Ng, A.W.T., Ng, A., Nicholson, J., Nicol, D., Nie, Y., Nielsen, G.P., Nielsen, M.M., Nik-Zainal, S., Noble, M.S., Nones, K., Northcott, P.A., Notta, F., O’Connor, B.D., O’Donnell, P., O’Donovan, M., O’Meara, S., O’Neill, B.P., O’Neill, J.R., Ocana, D., Ochoa, A., Oesper, L., Ogden, C., Ohdan, H., Ohi, K., Ohno-Machado, L., Oien, K.A., Ojesina, A.I., Ojima, H., Okusaka, T., Omberg, L., Ong, C.K., Ossowski, S., Ott, G., Ouellette, B.F.F., P’ng, C., Paczkowska, M., Paiella, S., Pairojkul, C., Pajic, M., Pan-Hammarström, Q., Papaemmanuil, E., Papatheodorou, I., Paramasivam, N., Park, J.W., Park, J.-W., Park, K., Park, K., Park, P.J., Parker, J.S., Parsons, S.L., Pass, H., Pasternack, D., Pastore, A., Patch, A.-M., Pauporté, I., Pea, A., Pearson, J.V., Pedamallu, C.S., Pedersen, J.S., Pederzoli, P., Peifer, M., Pennell, N.A., Perou, C.M., Perry, M.D., Petersen, G.M., Peto, M., Petrelli, N., Petryszak, R., Pfister, S.M., Phillips, M., Pich, O., Pickett, H.A., Pihl, T.D., Pillay, N., Pinder, S., Pinese, M., Pinho, A.V., Pitkänen, E., Pivot, X., Piñeiro-Yáñez, E., Planko, L., Plass, C., Polak, P., Pons, T., Popescu, I., Potapova, O., Prasad, A., Preston, S.R., Prinz, M., Pritchard, A.L., Prokopec, S.D., Provenzano, E., Puente, X.S., Puig, S., Puiggròs, M., Pulido-Tamayo, S., Pupo, G.M., Purdie, C.A., Quinn, M.C., Rabionet, R., Rader, J.S., Radlwimmer, B., Radovic, P., Raeder, B., Raine, K.M., Ramakrishna, M., Ramakrishnan, K., Ramalingam, S., Raphael, B.J., Rathmell, W.K., Rausch, T., Reifenberger, G., Reimand, J., Reis-Filho, J., Reuter, V., Reyes-Salazar, I., Reyna, M.A., Reynolds, S.M., Rheinbay, E., Riazalhosseini, Y., Richardson, A.L., Richter, J., Ringel, M., Ringnér, M., Rino, Y., Rippe, K., Roach, J., Roberts, L.R., Roberts, N.D., Roberts, S.A., Robertson, A.G., Robertson, A.J., Rodriguez, J.B., Rodriguez-Martin, B., Rodríguez-González, F.G., Roehrl, M.H.A., Rohde, M., Rokutan, H., Romieu, G., Rooman, I., Roques, T., Rosebrock, D., Rosenberg, M., Rosenstiel, P.C., Rosenwald, A., Rowe, E.W., Royo, R., Rozen, S.G., Rubanova, Y., Rubin, M.A., Rubio-Perez, C., Rudneva, V.A., Rusev, B.C., Ruzzenente, A., Rätsch, G., Sabarinathan, R., Sabelnykova, V.Y., Sadeghi, S., Sahinalp, S.C., Saini, N., Saito-Adachi, M., Saksena, G., Salcedo, A., Salgado, R., Salichos, L., Sallari, R., Saller, C., Salvia, R., Sam, M., Samra, J.S., Sanchez-Vega, F., Sander, C., Sanders, G., Sarin, R., Sarrafi, I., Sasaki-Oku, A., Sauer, T., Sauter, G., Saw, R.P.M., Scar-doni, M., Scarlett, C.J., Scarpa, A., Scelo, G., Schadendorf, D., Schein, J.E., Schilhabel, M.B., Schlesner, M., Schlomm, T., Schmidt, H.K., Schramm, S.-J., Schreiber, S., Schultz, N., Schumacher, S.E., Schwarz, R.F., Scolyer, R.A., Scott, D., Scully, R., Seethala, R., Segre, A.V., Selander, I., Semple, C.A., Senbabaoglu, Y., Sengupta, S., Sereni, E., Serra, S., Sgroi, D.C., Shackleton, M., Shah, N.C., Shahabi, S., Shang, C.A., Shang, P., Shapira, O., Shelton, T., Shen, C., Shen, H., Shepherd, R., Shi, R., Shi, Y., Shiah, Y.-J., Shibata, T., Shih, J., Shimizu, E., Shimizu, K., Shin, S.J., Shiraishi, Y., Shmaya, T., Shmulevich, I., Shorser, S.I., Short, C., Shrestha, R., Shringarpure, S.S., Shriver, C., Shuai, S., Sidiropoulos, N., Siebert, R., Sieuwerts, A.M., Sieverling, L., Signoretti, S., Sikora, K.O., Simbolo, M., Simon, R., Simons, J.V., Simpson, J.T., Simpson, P.T., Singer, S., Sinnott-Armstrong, N., Sipahimalani, P., Skelly, T.J., Smid, M., Smith, J., Smith-McCune, K., Socci, N.D., Sofia, H.J., Soloway, M.G., Song, L., Sood, A.K., Sothi, S., Sotiriou, C., Soulette, C.M., Span, P.N., Spellman, P.T., Sperandio, N., Spillane, A.J., Spiro, O., Spring, J., Staaf, J., Stadler, P.F., Staib, P., Stark, S.G., Stebbings, L., Stefánsson, A., Stegle, O., Stein, L.D., Stenhouse, A., Stewart, C., Stilgenbauer, S., Stobbe, M.D., Stratton, M.R., Stretch, J.R., Struck, A.J., Stuart, J.M., Stunnenberg, H.G., Su, H., Su, X., Sun, R.X., Sungalee, S., Susak, H., Suzuki, A., Sweep, F., Szczepanowski, M., Sültmann, H., Yugawa, T., Tam, A., Tamborero, D., Tan, B.K.T., Tan, D., Tan, P., Tanaka, H., Taniguchi, H., Tan-skanen, T.J., Tarabichi, M., Tarnuzzer, R., Tarpey, P., Taschuk, M.L., Tatsuno, K., Tavaré, S., Taylor, D.F., Taylor-Weiner, A., Teague, J.W., Teh, B.T., Tembe, V., Temes, J., Thai, K., Thayer, S.P., Thiessen, N., Thomas, G., Thomas, S., Thompson, A., Thompson, A.M., Thompson, J.F.F., Thompson, R.H., Thorne, H., Thorne, L.B., Thorogood, A., Tiao, G., Tijanic, N., Timms, L.E., Tirabosco, R., Tojo, M., Tommasi, S., Toon, C.W., Toprak, U.H., Torrents, D., Tortora, G., Tost, J., Totoki, Y., Townend, D., Traficante, N., Treilleux, I., Trotta, J.-R., Trümper, L.H.P., Tsao, M., Tsunoda, T., Tubio, J.M.C., Tucker, O., Turkington, R., Turner, D.J., Tutt, A., Ueno, M., Ueno, N.T., Umbricht, C., Umer, H.M., Underwood, T.J., Urban, L., Urushidate, T., Ushiku, T., Uusküla-Reimand, L., Valencia, A., Van Den Berg, D.J., Van Laere, S., Van Loo, P., Van Meir, E.G., Eynden, G.G., Kwast, T., Vasudev, N., Vazquez, M., Vedururu, R., Veluvolu, U., Vembu, S., Verbeke, L.P.C., Vermeulen, P., Verrill, C., Viari, A., Vicente, D., Vicentini, C., VijayRaghavan, K., Viksna, J., Vilain, R.E., Villasante, I., Vincent-Salomon, A., Visakorpi, T., Voet, D., Vyas, P., Vázquez-García, I., Waddell, N.M., Waddell, N., Wadelius, C., Wadi, L., Wagener, R., Wala, J.A., Wang, J., Wang, J., Wang, L., Wang, Q., Wang, W., Wang, Y., Wang, Z., Waring, P.M., Warnatz, H.-J., Warrell, J., Warren, A.Y., Waszak, S.M., Wedge, D.C., Weichenhan, D., Weinberger, P., Weinstein, J.N., Weischenfeldt, J., Weisenberger, D.J., Welch, I., Wendl, M.C., Werner, J., Whalley, J.P., Wheeler, D.A., Whitaker, H.C., Wigle, D., Wilkerson, M.D., Williams, A., Wilmott, J.S., Wilson, G.W., Wilson, J.M., Wilson, R.K., Winterhoff, B., Wintersinger, J.A., Wiznerowicz, M., Wolf, S., Wong, B.H., Wong, T., Wong, W., Woo, Y., Wood, S., Wouters, B.G., Wright, A.J., Wright, D.W., Wright, M.H., Wu, C.-L., Wu, D.-Y., Wu, G., Wu, J., Wu, K., Wu, Y., Wu, Z., Xi, L., Xia, T., Xiang, Q., Xiao, X., Xing, R., Xiong, H., Xu, Q., Xu, Y., Xue, H., Yachida, S., Yakneen, S., Yamaguchi, R., Yamaguchi, T.N., Yamamoto, M., Yamamoto, S., Yamaue, H., Yang, F., Yang, H., Yang, J.Y., Yang, L., Yang, L., Yang, S., Yang, T.-P., Yang, Y., Yao, X., Yaspo, M.-L., Yates, L., Yau, C., Ye, C., Ye, K., Yellapantula, V.D., Yoon, C.J., Yoon, S.-S., Yousif, F., Yu, J., Yu, K., Yu, W., Yu, Y., Yuan, K., Yuan, Y., Yuen, D., Yung, C.K., Zaikova, O., Zamora, J., Zapatka, M., Zenklusen, J.C., Zenz, T., Zeps, N., Zhang, C.-Z., Zhang, F., Zhang, H., Zhang, H., Zhang, H., Zhang, J., Zhang, J., Zhang, J., Zhang, X., Zhang, X., Zhang, Y., Zhang, Z., Zhao, Z., Zheng, L., Zheng, X., Zhou, W., Zhou, Y., Zhu, B., Zhu, H., Zhu, J., Zhu, S., Zou, L., Zou, X., deFazio, A., As, N., Deurzen, C.H.M., Vijver, M.J., Veer, L., Mering, C.: The repertoire of mutational signatures in human cancer. Nature 578(7793), 94–101 (2020) 10.1038/s41586-020-1943-3

[14] Kaplanis, J., Ide, B., Sanghvi, R., Neville, M., Danecek, P., Coorens, T., Prigmore, E., Short, P., Gallone, G., McRae, J., Moutsianas, L., Odhams, C., Carmichael, J., Barnicoat, A., Firth, H., O’Brien, P., Rahbari, R., Hurles, M.: Genetic and chemotherapeutic influences on germline hypermutation. Nature 605(7910), 503–508 (2022) 10.1038/s41586-022-04712-2

[15] Supek, F., Lehner, B.: Differential dna mismatch repair underlies mutation rate variation across the human genome. Nature 521(7550), 81–84 (2015) 10.1038/nature14173

[16] Harris, K., Pritchard, J.K.: Rapid evolution of the human mutation spectrum. eLife 6 (2017) 10.7554/elife.24284

[17] Mathieson, I., Reich, D.: Differences in the rare variant spectrum among human populations. PLOS Genetics 13(2), 1006581 (2017) 10.1371/journal.pgen.1006581

